# Highly Multiplexed Proteomic Analysis of HCMV-Infected Dendritic Cells Reveals Global Manipulation of Adaptive Immunity and Host Restriction of Viral Replication

**DOI:** 10.1101/2024.04.30.591855

**Authors:** Lauren E. Kerr-Jones, Lior Soday, Nia Cwyfan Hughes, Xinyue Wang, Leah H. Hunter, Robin Antrobus, Kelly L. Miners, Kristin Ladell, David A. Price, Ceri A. Fielding, Eddie C. Y. Wang, Michael P. Weekes, Richard J. Stanton

**Affiliations:** Division of Infection and Immunity, School of Medicine, Cardiff University, Cardiff, CF14 4XN; Cambridge Institute for Medical Research, University of Cambridge, Hills Road, Cambridge CB2 0XY, UK; Department of Medicine, University of Cambridge, Hills Road, Cambridge CB2 0XY, UK

**Author notes:** These authors contributed equally (Lauren E. Kerr-Jones, Lior Soday). Senior authors (Michael P. Weekes, Richard J. Stanton).

## Abstract

Human cytomegalovirus (HCMV) is a clinically significant herpesvirus and a paradigm for pathogen-mediated immune-evasion. Its broad tropism includes antigen-presenting cells such as dendritic cells (DCs), which may partly explain a unique, dramatic imprint on host immunity that occurs following lifelong carriage. Despite this breadth of infection, most studies use fibroblasts as a model. We therefore developed systems to isolate pure populations of DCs infected with wild-type HCMV, before applying quantitative temporal proteomic technologies to systematically characterise the virus:DC interaction within cells and at the cell surface. This comprehensive dataset quantifying almost 9,000 proteins throughout the infection timecourse revealed multiple DC-specific viral:host effects, including key impacts on innate, intrinsic, and adaptive immunity. These effects included observations that APOBEC3A is downregulated in infected cells and restricts HCMV infection in *ex vivo* DCs, delaying the progression of lytic infection, and that cell surface ICOS-Ligand was downregulated by the viral genes US16 and US20, inhibiting the induction of adaptive immunity.

## Introduction

Human cytomegalovirus (HCMV) establishes lifelong infection despite inducing robust humoral and cellular adaptive immunity. It is one of the most common infectious causes of complications in the immunocompromised, particularly transplant recipients, and the most common infectious cause of congenital malformation. Combined with the high seroprevalence and global distribution of HCMV, it creates a significant financial burden on healthcare systems worldwide^1^.

HCMV has the biggest genome of any human virus, and induces one of the largest immune responses seen to any pathogen^2^. However, over its long co-evolution with its host, it has evolved to dedicate a substantial proportion of its 235kb genome to manipulating human immunity. As a result, despite broad and persistent antiviral responses involving both innate and adaptive immunity, HCMV persists lifelong, and immunity is insufficient to prevent repeated superinfection. Studies of this phenomenon have resulted in the virus becoming a paradigm for pathogen-mediated immune evasion, informing on the underlying functions of host immunity^3^, revealing immunological mechanisms that can be exploited to enhance vaccine and immunotherapeutic strategies against heterologous infections^4^ and tumours^5^, as well as providing some explanations for the unique lifelong remodelling of human immune system function that occurs following infection^6^. To date, these studies have overwhelmingly used infection of fibroblasts as a model cell system, and have focused on virus evasion of antiviral restriction factors, and NK-, B- and T-cell based immunity^7^. Yet *in vivo,* HCMV also infects antigen presenting cells (APCs), such as dendritic cells (DCs)^8^. DCs link innate and intrinsic immunity by presenting antigens to B- and T-cells, inducing their activation and proliferation^9^. However, despite a presumably critical requirement to evade APC-dependent immunity, HCMV infection of DCs has not been systematically studied.

Laboratory strains of HCMV that have been extensively passaged in permissive cell lines such as fibroblasts have been widely used in research, however these have accumulated multiple mutations^10^. Mutations are rapidly selected even in early passage clinical isolates, with RL13 mutations detectable as early as 3 passages, UL128-UL131A demonstrating mutations as early as 15 passages, and mutations in the UL/b’ region of the genome being selected after further growth *in vitro*^11^. Many of these genes have key roles in immune-regulation, latency, and tropism, and their mutation has major phenotypic effects on the virus. Importantly, although HCMV has broad tropism for multiple cell-types *in vivo*, including myeloid, endothelial, and epithelial cells, partial or complete loss of UL128L *in vitro* results in an inability to infect cells other than fibroblasts^11–13^. It also causes a switch from a form of virus spread that is almost completely cell-associated to one that is dominated by cell-free virion dissemination^14–16^. As a result, compared to virus found *in vivo*, laboratory strains can exhibit significant differences in immunological control, tropism, and mechanisms of spread^10^.

To address this issue, we generated a bacterial artificial chromosome (BAC) clone of a minimally passaged HCMV strain (Merlin) which had also been sequenced from patient material prior to isolation. We then repaired all *in vitro* acquired mutations, and engineered the BAC such that expression of the genes that mutate most rapidly during passage (RL13 and UL128L) could be repressed during viral replication in a cell line expressing the tetracycline repressor (TetR)^17^. This enabled a virus containing the complete repertoire of HCMV genes (hereafter referred to as wildtype HCMV (WT-HCMV) to be passaged *in vitro* without mutations being selected^15^. In cells lacking TetR, Merlin expresses all genes, and mimics clinical isolates by spreading predominantly through direct cell-cell contact, and efficiently infecting epithelial, endothelial and myeloid cells^16^.

To facilitate analysis of WT-HCMV infection in APCs, in the present study we have developed technologies permitting cell-associated WT-HCMV to be transferred to DCs from a donor cell, then established novel systems to enrich populations of infected or bystander DCs. Our ‘Quantitative Temporal Viromics’ (QTV) technology^18^ was then used to analyse proteome-wide changes within both whole-cell and plasma membrane proteins, across an infectious time course. This provides the first unbiased proteome-wide view of how HCMV infection alters DCs. It revealed that >1,100 proteins are modulated by infection, many of which were not quantified in previous analyses of HCMV infection in fibroblasts yet play key roles in the priming of adaptive immunity. This viral modulation impacts the development of CD4+ T-cell-mediated protective immunity, in part due to viral downregulation of the inducible costimulatory (ICOS) ligand (ICOSLG), mediated by US16 and US20. The data also revealed that primary DCs can limit productive virus infection via APOBEC3A (A3A), which inhibits genome replication and virion maturation^19^. Overall, this establishes new systems enabling dissection of WT HCMV infection in clinically relevant cell types, demonstrates that HCMV infection in DCs differs substantially from more standard *in vitro* models, defines novel ways in which HCMV manipulates host immunity, and identifies DCs as possessing unique antiviral systems that limit the impact of infection.

## Results

### Straightforward enrichment of primary DCs infected with WT-HCMV following co-culture

Although WT-HCMV transmits very efficiently from cell-to-cell, for example when infected donor cells are mixed with uninfected targets, subsequent analyses have hitherto been limited to single-cell based technologies because of the resulting mixed population^16^. Fluorescence-activated cell sorting (FACS) can resolve this problem but is not available routinely due to the specialised facilities required for sorting cells infected with live virus. We therefore developed a universally applicable system for purification of infected target cells in a standard BSL2 laboratory (**Figure 1A**), including: (1) a WT-HCMV strain that rapidly expresses rat CD2 (HCMV-rCD2) on the surface of infected cells. Here, rat CD2 has a truncated C-terminus to prevent signalling and is fused via a P2A peptide to the immediate-early viral protein UL36. (2) a telomerase immortalised fibroblast cell line (HFFF-His) expressing a 6xHis-tagged cell surface mCherry, enabling the depletion of infected donor cells. HCMV-rCD2 containing a complete wildtype genome was grown in fibroblasts expressing TetR to repress the expression of RL13 and UL128L^17^. This system prevents the selection of mutants^15^ and enables the release of high-titre cell-free virus^16^ that is then used to infect HFFF-His at high MOI. Since all viral genes including UL128L are now expressed, the virus becomes cell-associated and transmits very efficiently through the cell-cell route into uninfected DCs. DCs are separated from HFFF-His cells via negative magnetic-activated cell sorting (MACS), then infected DCs separated from bystander DCs in a second MACS step using anti-rat CD2 (**Figure 1A**). For the purposes of comparison, and to generate a biologically independent replicate for proteomic analysis, a system involving FACS-based separation was also employed. Here, DCs were co-cultured with HFFF-His cells infected with HCMV expressing GFP (HCMV-UL36-GFP), then DCs separated into GFP positive and negative populations by FACS. The purity of DC populations using both MACS and FACS-based systems was >95%, however the MACS-based method gave superior recovery (82% vs 50%).

**Figure 1.**
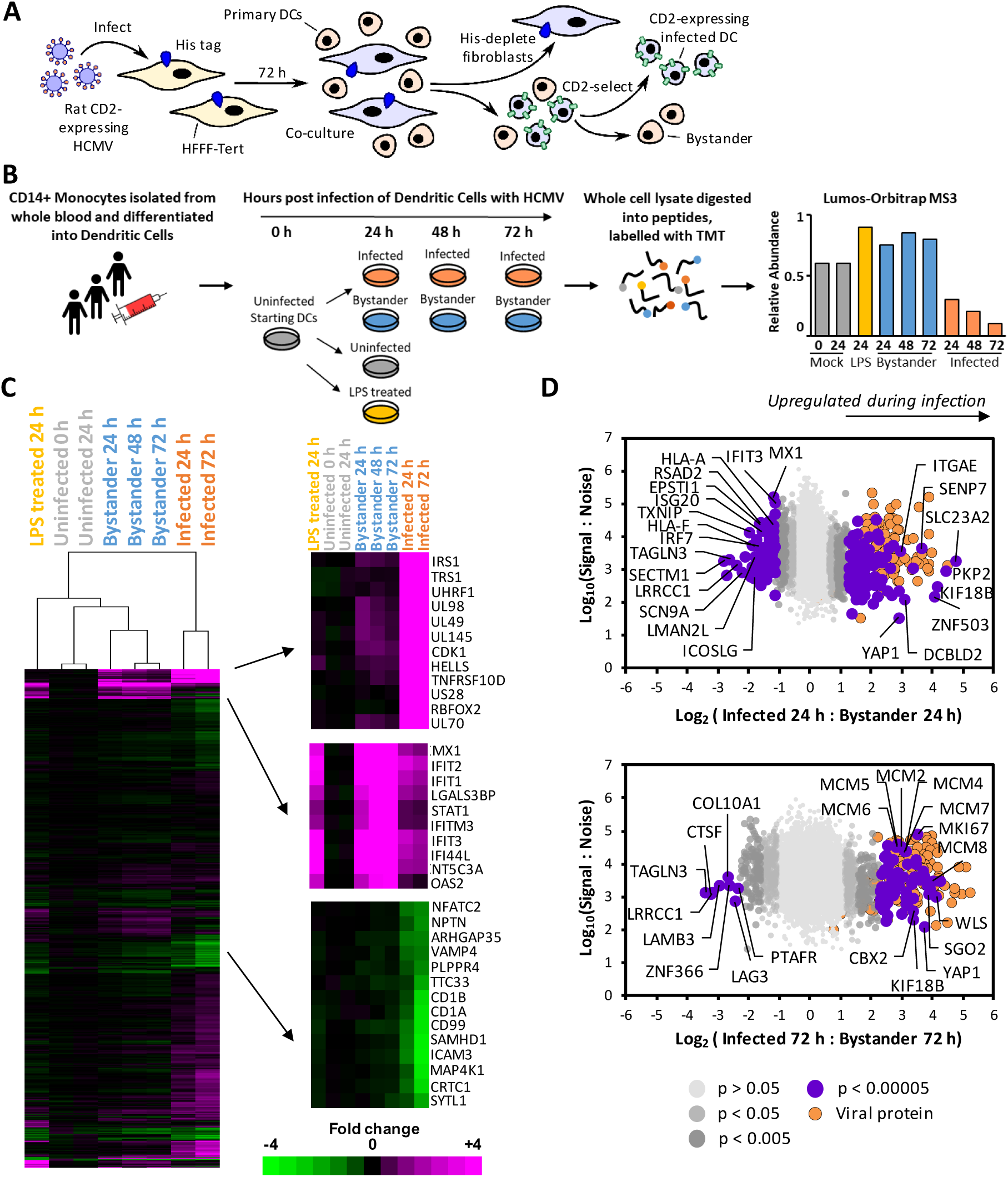
Quantitative temporal viromic analysis of whole cell lysates. (A) Schematic of methodology employed to enrich bystander and infected DCs. (B) Schematic of proteomic workflow. Infected and bystander DCs were derived as described in the main text, and compared to uninfected DCs, DCs uninfected and cultured for 24 h and DCs stimulated for 24h with lipopolysaccharide (LPS) (n=2). Data from 48h post-infection was examined in replicate WCL1 only due to the lower recovery of cells in WCL2. For this reason, averaged data from the 24h and 72h timepoints are analysed throughout the subsequent text apart from where indicated. All data are shown in the ‘plotter’ in **Table S2**. (C) Hierarchical cluster analysis of all 8844 proteins quantified in either WCL1 or WCL2. The fold change is shown as compared to the average S:N of the two uninfected time points (0 and 24 hpi). For proteins quantified in both WCL1 and WCL2, an average fold change was taken for each sample. An enlargement of three subclusters is shown on the right, including multiple proteins that were substantially up- or down-regulated during infection (top and bottom), and proteins that were predominantly upregulated in the bystander samples (middle), many of which are known to be interferon-stimulated genes. (D) Scatterplots of all 8844 proteins quantified in either WCL1 or WCL2, at 24 (top) or 72 hpi (bottom). The fold change and total S:N was averaged for proteins quantified in both experiments. Benjamini-Hochberg adjusted Significance A values were used to estimate p-values (see Methods).

### Multiplexed proteomic comparison of whole cell changes in HCMV-infected and bystander primary DCs

For immune cells such as APCs, many of the key interactions that define the outcome of infection occur via proteins expressed at the plasma membrane. We therefore applied our ‘Quantitative Temporal Viromic’ (QTV) technology^18^, to infected and bystander DCs across a timecourse of infection (**Figure 1B**). QTV employs TMT labelling and MS3 mass spectrometry to globally quantify virus induced changes both within and at the surface of infected cells. This technology can therefore directly measure infection-related changes at transcriptional, translational and post-translational levels. Critically for HCMV infection, this captures key mechanisms of immune evasion including protein relocalisation, targeted degradation, and retention from the plasma membrane in intracellular stores^3,7,20,21^.

Two biological replicates were performed for proteomic analysis, one using the MACS-based enrichment (whole cell lysate 1 (WCL1)) and plasma membrane 1 (PM1)) and a separate donor using FACS (WCL2 and PM2). In whole cell lysates, an overall total of 8717 human and 127 viral proteins were quantified in either replicate (7474 and 96 proteins respectively in both), providing a global view of changes in protein expression during infection (**Figure S1A**). Protein changes were largely independent of the method of separation or donor employed, with strong correlation between MACS- and FACS-enriched samples both early and late during infection (**Figure S1B**). Uninfected and bystander samples clustered separately from early / late infection time points in both replicates and in combined data. Overall, 486 proteins were downregulated >2-fold and 655 proteins upregulated in infected compared to bystander cells at either 24 or 72 h post-infection (hpi), with the greatest number of protein changes occurring late during infection (**Figures 1C-D, S1A, S1C-D**). Interestingly, 283 proteins were upregulated >2-fold in bystander cells compared to uninfected cells, with proteins changing in a similar manner to control lipopolysaccharide (LPS)-stimulated DCs (**Figure S2A-B**). Multiple interferon-stimulated genes were upregulated in both cases, suggesting that the cause was likely to be cytokine release from infected DCs and donor HFFFs, and emphasising the utility of bystander controls for comparison with infected cells to distinguish true infection-related protein changes (**Figure S2C-D, Table S1A-B**). All data are shown in **Table S2**, in which the worksheet ‘‘Plotter’’ enables interactive generation of temporal graphs of the expression of each of the human or viral proteins quantified.

### Regulation of multiple molecules with key roles in immunity

We applied DAVID software to determine which pathways were enriched amongst proteins downregulated >2-fold in infected compared to bystander cells^22^. The results suggested that multiple proteins important in all forms of intrinsic, innate and adaptive immunity are downregulated throughout infection. These included components of viral sensing pathways, interferon signalling pathways, cell surface proteins important in adhesion and antigen presentation, and factors with roles in cytokine signalling (**Figure S3A, Table S3A**). Proteins upregulated >2-fold were enriched in factors important in DNA replication, chromosome maintenance and cell cycle regulation (**Figure S3B, Table S3B**).

Most antiviral factors are induced by interferon or infection, and a subset are specifically inhibited, downregulated or degraded by viral proteins^23^. To identify candidate factors that have the potential to play antiviral roles during HCMV infection in DCs, and are therefore antagonised by the virus, we performed a combined search for proteins that were induced in bystander cells but comparatively downregulated in infected cells (**Table S3C**). Enrichment analysis revealed modulation of an array of at least 45 proteins important in antiviral defence as well as proteins with roles in production of and signalling by interferons α, β and γ and tumour necrosis factor α. These proteins included interferon regulated interferon regulated factors (IRF) 1, 7, 8 and 9; Janus-associated kinase 1 (JAK1), signal transducer and activator of transcription 1 (STAT1), and multiple proteins with known roles in intrinsic immunity against HCMV and other viruses (**Figure 2, Table S3C**).

**Figure 2.**
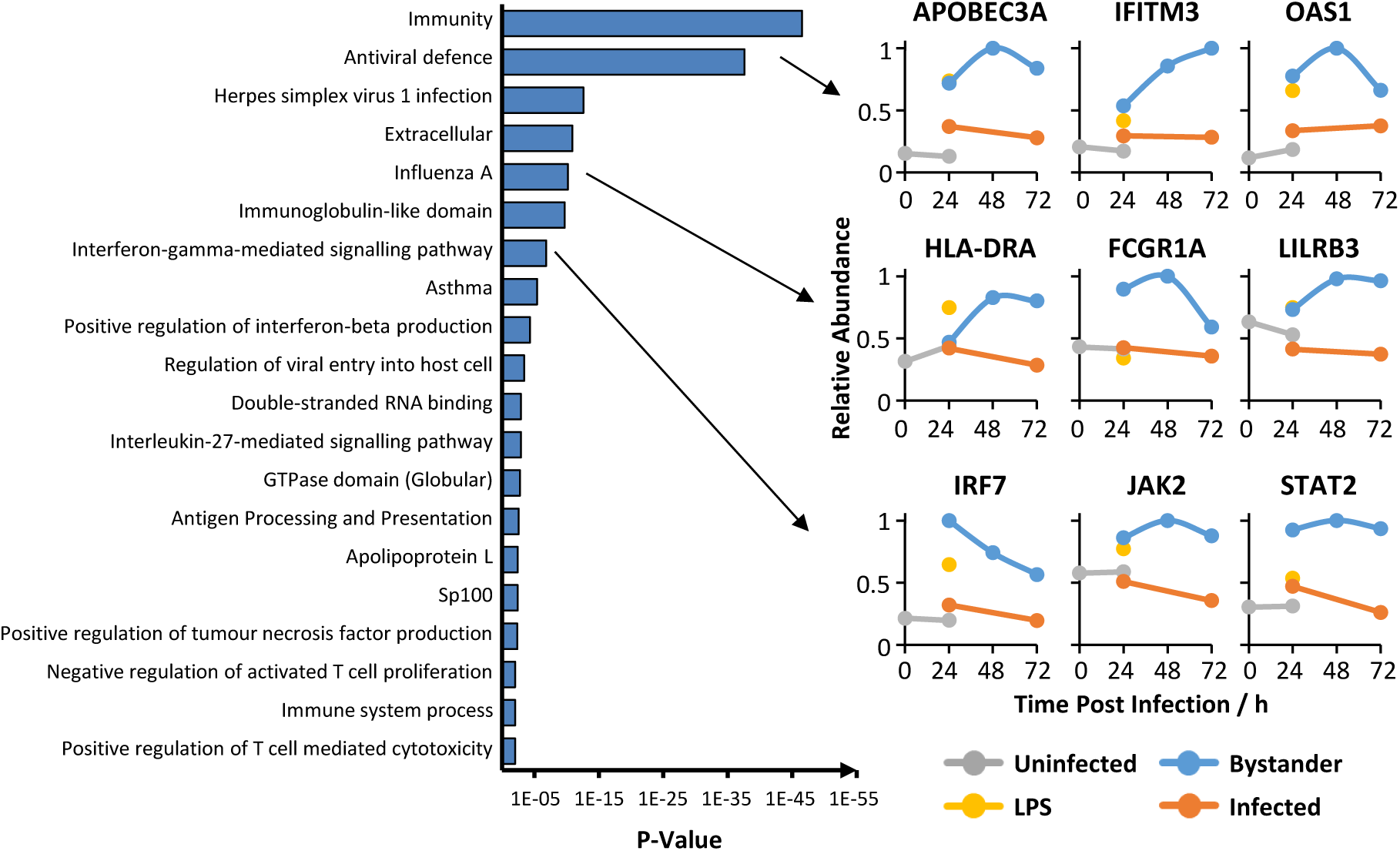
Regulation of multiple immune pathways during HCMV infection of DCs. Functional enrichment analysis of 307 proteins induced >1.5 fold in bystander cells but comparatively downregulated >1.5 fold in infected cells, at either 24 or 72h post infection. Displayed are significantly enriched clusters (Benjamini corrected p < 0.01) with example protein profiles. Full data for clusters enriched at p < 0.05 are shown in **Table S3C**.

### Comparative analysis of protein regulation in DCs and HFFFs

To analyse the effects of HCMV infection in different cell types systematically, we compared data from the present study with our prior temporal analysis of infection in HFFFs. 7655 proteins were quantified in both cell types, with 1060 and 1097 quantified only in DCs or HFFFs respectively. Whilst many of the more substantial protein changes in DCs also changed in the same direction in HFFFs, albeit with differences in absolute magnitude, overall the correlation between changes in both cell types was modest (r^2^ = 0.10, Figure 3A). Interestingly, correlation was much better for the subset of proteins we previously defined as targeted for degradation from early times of HCMV infection in HFFFs^21^. For 28/35 proteins degraded with the highest confidence and quantified in DCs, r^2^ was 0.47 (**Figure 3A, Table S4A**), and for 108/133 proteins degraded with medium confidence and quantified in DCs, r^2^ was 0.23 (**Table S4A**). This suggests that the degradation machinery for many proteins targeted in HFFFs is preserved in DCs, and that many of the protein changes that do not involve HCMV-directed protein degradation may reflect a cell-type specific response to infection.

**Figure 3.**
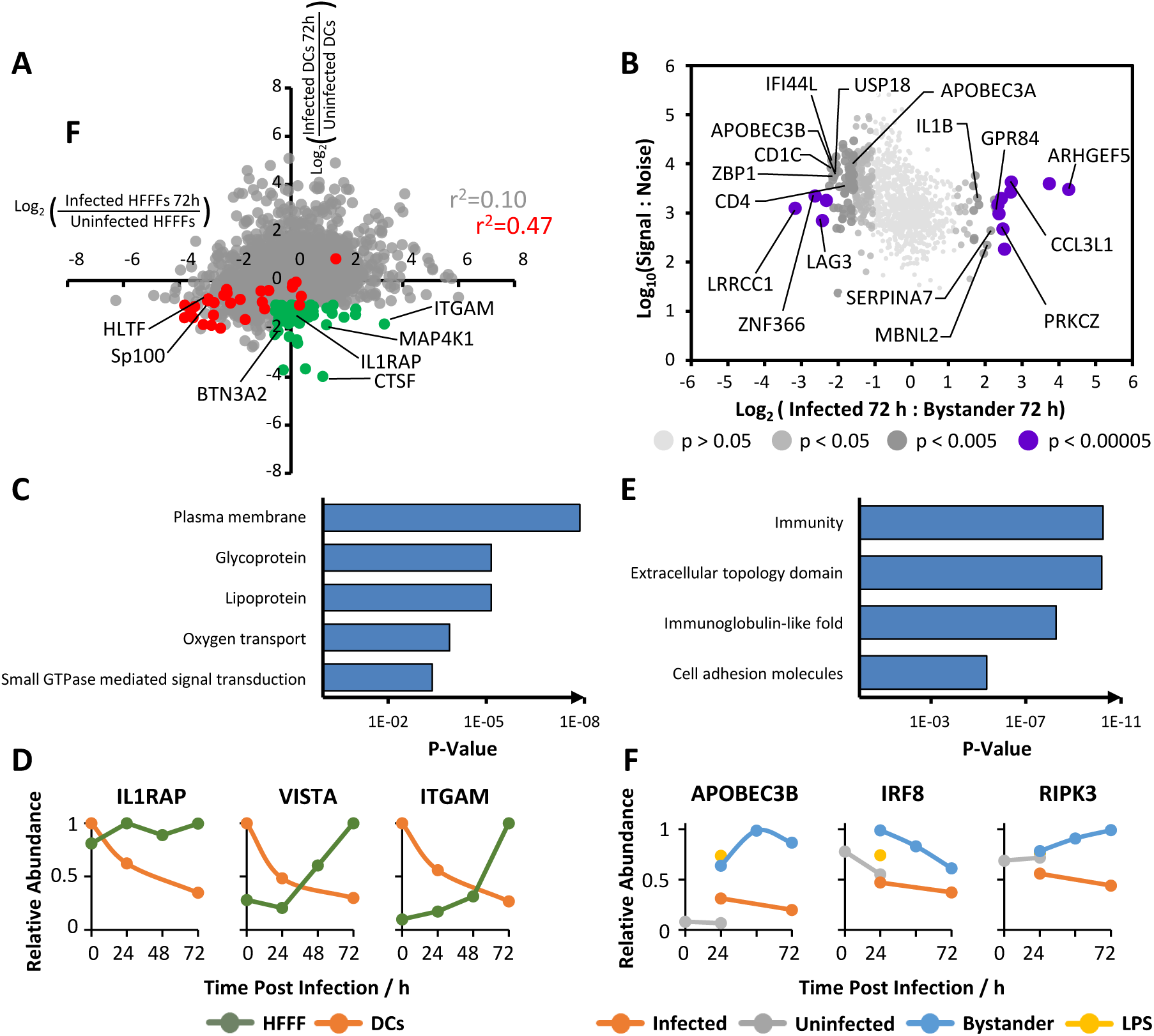
Unique regulation of a subset of human proteins in DCs but not in HFFFs. (A) Comparison of protein regulation by HCMV in DCs with HFFFs, for 7655 proteins quantified in both cell types. For DCs, fold changes were calculated for proteins quantified in either experiment WCL1 or WCL2, with average fold change shown where quantified in both. For HFFFs, fold changes were calculated at 24, 48 and 72h for proteins quantified in any of experiments WCL1, WCL2 or WCL3^18,24^, with average fold change shown where quantified in more than one experiment. Green dots indicate proteins that were downregulated >2-fold in DCs at 72h, but downregulated <1.5-fold in HFFFs at all of 24, 48, and 72 h. Red dots indicate 28/35 proteins that are degraded with high confidence early during HCMV infection in HFFFs^21^, and quantified in DCs. Of note, a few of these proteins (e.g. GLG1, MORC3) were downregulated early during infection but upregulated again by 72 h. A full list of proteins degraded with medium and high confidence and their changes in HFFFs and DCs is shown in **Table S4A**. (B) Scatterplot of 1060 proteins only quantified in DCs but not in HFFFs, at 72 hpi. The fold change and total S:N was averaged for proteins quantified in both experiments. Benjamini-Hochberg adjusted Significance A values were used to estimate p-values, as calculated for **Figure D**. (C) Functional enrichment analysis of the 76 proteins downregulated >2-fold in DCs at 24 and/or 72h (the latter highlighted green in Figure 3A), but downregulated <1.5-fold in HFFFs at all of 24, 48 and 72h. Displayed are significantly enriched clusters (Benjamini corrected p < 0.01) with example protein profiles. Full data for clusters enriched at p < 0.05 are shown in **Table S4A**. (D) Examples of proteins downregulated only in DCs (highlighted in green in **Figure 3A**). (E) Functional enrichment analysis of 219 proteins only quantified in DCs and downregulated >2 fold in comparison to bystander cells at either 24 or 72h post infection. Displayed are significantly enriched clusters (Benjamini corrected p < 0.01) with example protein profiles. Full data for clusters enriched at p < 0.05 are shown in **Table S4B**. (F) Examples of proteins quantified and downregulated only in DCs (highlighted in red in **Figure 3A**).

Of particular interest were proteins that were either quantified in both cell types but only downregulated by HCMV in DCs (**Figure 3A**), or only quantified and downregulated in DCs (**Figure 3B**), since these might represent cell type-specific mechanisms of antiviral control that are regulated by HCMV. The former category included multiple cell surface receptors such as V-type immunoglobulin domain-containing suppressor of T-cell activation (VISTA, C10orf54), IL1 receptor-associated protein (IL1RAP) and integrin alpha M (ITGAM) (**Figures 3C-D, Table S4A**). The latter category also included multiple cell adhesion molecules and lectins, necroptosis mediator receptor interacting serine/threonine kinase 3 (RIPK3) and two cytidine deaminases, APOBEC3A (A3A) and -3B (A3B) (**Figures 3E-F, Table S4B**).

### Restriction of the HCMV replication cycle in DCs

Productive HCMV infection is traditionally separated into four phases of gene expression: immediate-early (IE), early (E), early-late (EL) and late (L), however this was extended to five temporal protein (Tp) classes in our prior QTV-based analysis^18^. During infection in HFFFs, Tp1 proteins are maximally expressed by 6-12h after infection and Tp5 proteins are expressed at the greatest levels from 72h, as new virions assemble. In DCs, there was a shift in the relative abundance of Tp1-Tp5 protein in comparison to HFFFs. Most strikingly, although proteins from earlier temporal classes were expressed at either equal or higher levels in DCs compared to HFFFs, Tp5 proteins were significantly reduced in abundance, with a similarly substantial reduction in Tp4 proteins that did not reach significance due to the limited number of proteins in this category.

To determine whether the expression of Tp5 proteins was simply delayed beyond 72 h in DCs, we engineered Merlin-strain HCMV to express UL36-P2A-mCherry as a marker of early-expressed proteins, and UL32-eGFP as a marker of Tp5 proteins. In infected HFFFs, expression of UL32 increased from 48 h, at which point virion production could be observed in the perinuclear viral assembly complex (VAC). A more complete VAC was visible from 72 h onwards in the majority of cells (**Figures 4C-D**). In contrast, in DCs, UL32 expression was not detectable until 96 h, and only a small proportion of infected cells progressed to the late stage of infection (**Figures 4E-F**). Thus, following cell-cell transmission of WT-HCMV, late-stage HCMV infection in DCs is delayed in comparison to HFFFs, and progression is restricted such that only a subpopulation of cells exhibit the full lytic cycle.

**Figure 4.**
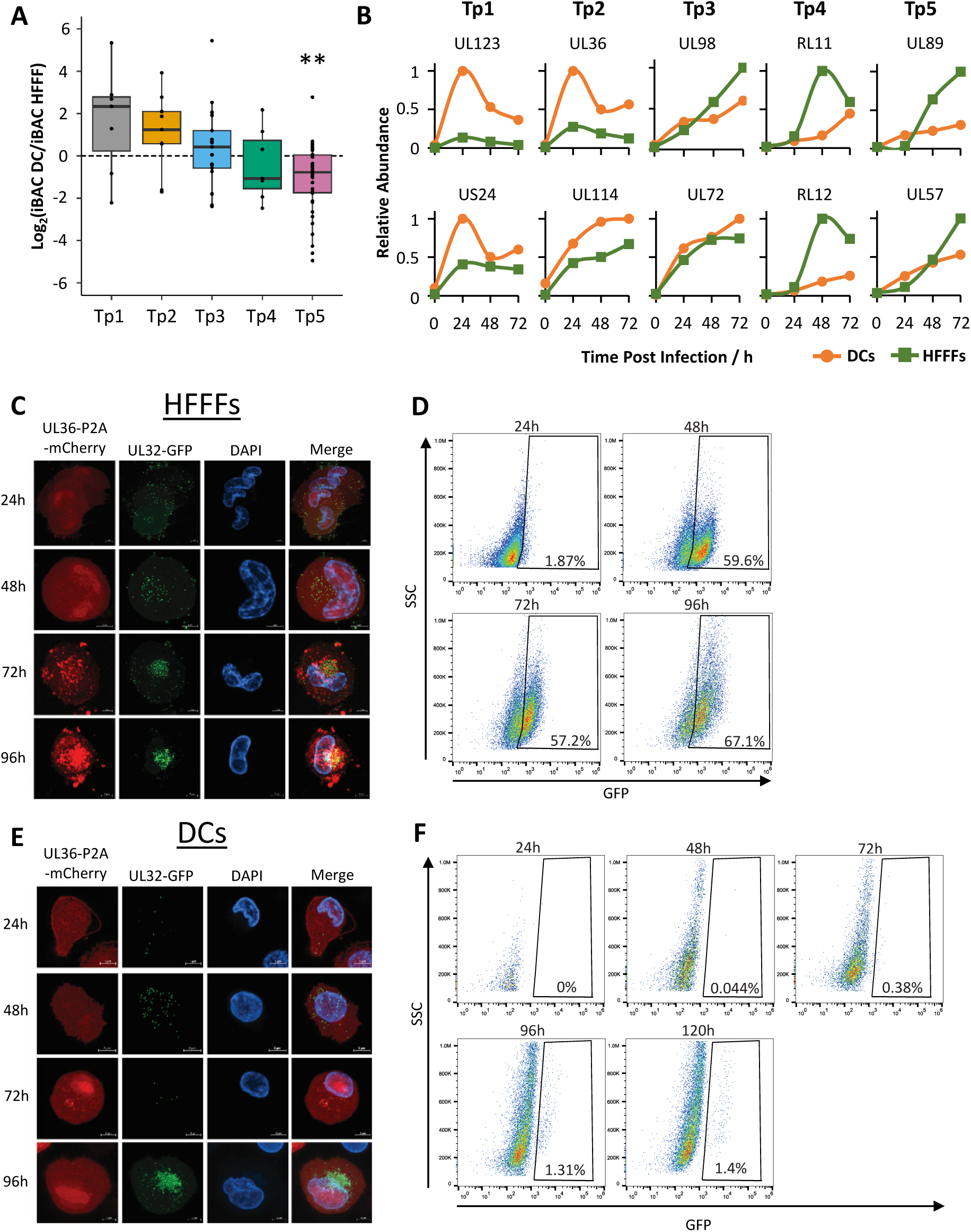
Restriction of the HCMV replication cycle in DCs. (A) Boxplot showing the relative abundance of HCMV proteins in DCs and HFFFs. iBAQ abundance values for viral proteins in DCs (see Methods) were compared to values previously generated for HFFFs^25^. Prior to calculation of the DC:HFFF iBAQ ratio, for each viral protein quantified in both cell types, iBAQ values were first normalised to the total abundance of these proteins. For example, the DC iBAQ value for UL123 was normalised to the total of DC iBAQ values for viral proteins quantified in both DCs and HFFFs. The boxplot shows 25^th^ and 75^th^ percentiles and median fold changes. Whiskers show the estimated range, and any data points beyond this range are considered outliers. Probabilities that the median log_2_ fold change was significantly different to 0 for each Tp class were calculated using a one-sample Wilcoxon signed rank test and adjusted for multiple hypothesis testing using the Benjamini-Hochberg method. **p < 0.01. (B) Example temporal graphs showing the relative abundance of individual viral proteins in DCs and HFFFs over time. To enable comparison between cell types, for DCs, the relative abundance for each viral protein was first adjusted using its DC:HFFF iBAQ ratio (shown in **Figure 4A** and **Table S5**), then all values for each protein were normalised to a maximum value of 1. (C-D) Expression of early and late HCMV proteins in HFFFs. HFFF were infected with HCMV expressing UL36-P2A-mCherry and UL32-eGFP for the indicated times, then either fixed and imaged by confocal microscopy (C), or trypsinised and analysed by flow cytometry (D). UL32 expression and VAC formation were visible from 48h onwards. (E-F) Expression of early and late HCMV proteins in DCs are delayed. DCs were infected with HCMV (as in (C)) by co-culture, then purified by MACS before being imaged by confocal microscopy (E) or flow cytometry (F). UL32 gene expression and VAC formation were not visible until 96 h, and then only in a small proportion of cells.

### A3A restricts HCMV genome replication in DCs

Transition to the late phase of the HCMV lytic cycle is dependent on viral DNA replication. Normal expression of Tp1 – Tp3 proteins in DCs with delayed Tp4 and delayed or absent Tp5 protein expression (**Figure 4**) suggested that viral DNA replication was inhibited. To identify human factors that might restrict HCMV replication in DCs but not HFFFs, we determined which proteins had known roles in antiviral defence against other viruses (**Figure 2, Table S3C**), yet were selectively antagonised (**Figure 3A**) or selectively expressed (**Figure 3B**) in DCs but not HFFFs (**Table S4**). Two obvious candidates were cytidine deaminases A3A and A3B, which have been reported to prevent the replication of other viruses (e.g. adeno-associated virus type 2 (AAV-2) and hepatitis B virus (HBV)) by promoting viral genome mutation during replication^19^.

To confirm that HCMV replication was restricted in primary DCs and also determine whether A3A played a role in this process, we knocked down A3A (**Table S8**) and quantified viral genomes across an infectious timecourse. In control HFFFs, HCMV genome replication was evident by 72 h as expected (**Figure 5A**). In DCs it was only possible to deplete A3A levels transiently using siRNA, nevertheless, in A3A-KD DCs genome levels increased much more substantially than in control cells (**Figure 5B**). To determine the impact of A3A-mediated restriction on the egress of *de novo* produced virions, DCs infected with HCMV-UL36-GFP were incubated on a monolayer of uninfected indicator HFFFs, and GFP fluorescence monitored over time. Although a substantial increase in GFP fluorescence indicative of viral lytic replication in the HFFFs did not occur until ∼6 days post-co-culture (9 days after the DCs were infected), virus transfer did eventually occur, and was significantly more efficient when A3A had been previously knocked down (**Figure 5C**). Thus, in DCs, virus genome replication and subsequent virus release is limited by A3A, but not prevented completely.

**Figure 5.**
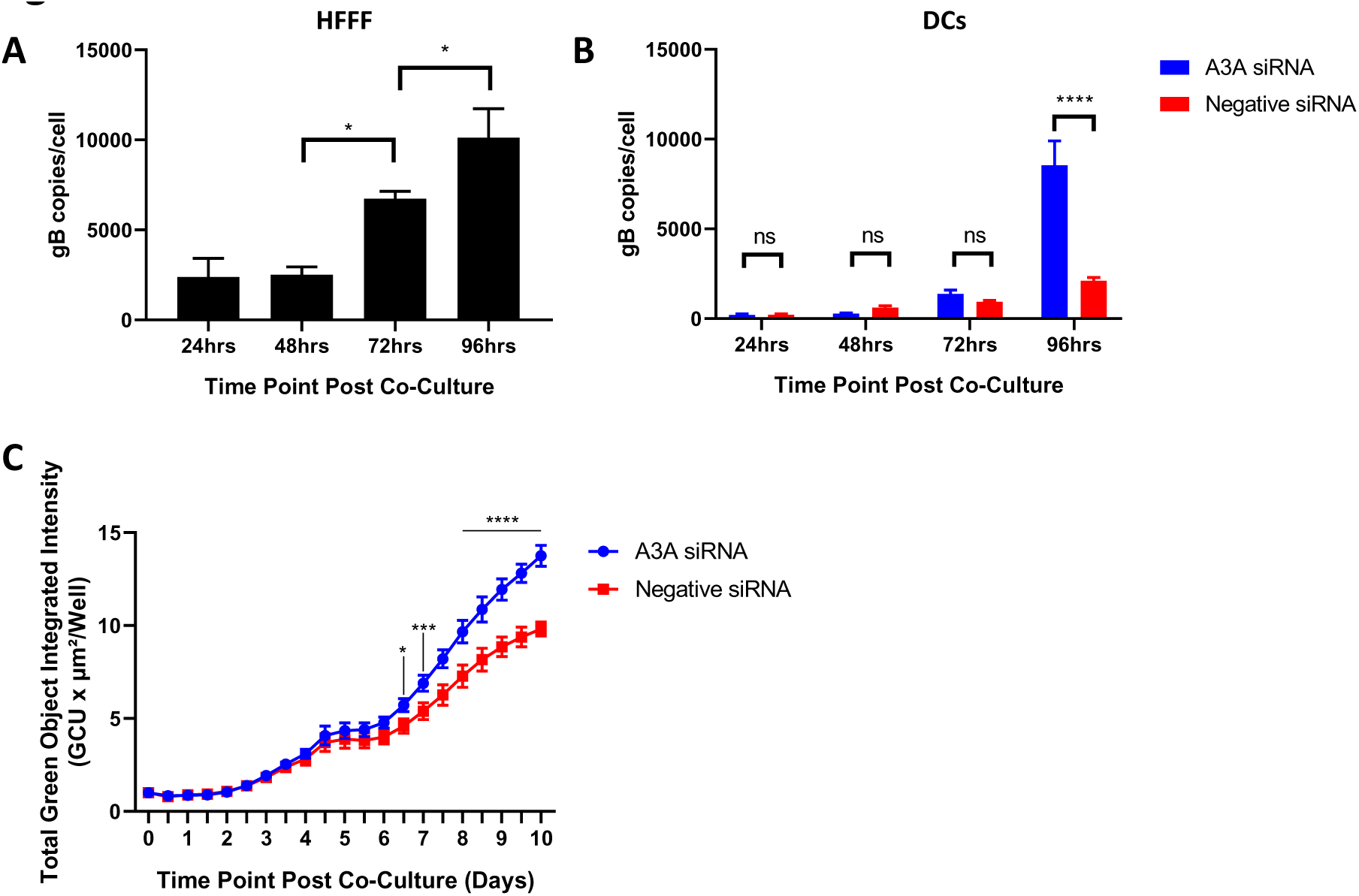
A3A restricts HCMV genome replication in DCs. (A) HCMV Genome copy numbers over time in HFFFs. HFFFs were infected by co-culture with HCMV-rCD2-infected HFFF-His cells (four uninfected HFFFs per infected HFFF), then newly infected cells purified by MACS at 24h. DNA was extracted from the cells at the indicated times and gB quantified relative to GAPDH using qPCR. *p < 0.05, by 1-way ANOVA (Tukey’s multiple comparisons test). (B) HCMV genome copy numbers over time in DCs. DCs were pre-treated with A3A-targeting or negative control siRNA, and infected 48 h later by co-culture with HCMV-rCD2-infected HFFF-His cells. Newly infected DCs were purified using MACS. DNA was extracted from the cells over a timecourse and gB quantified relative to GAPDH using qPCR. ****p < 0.0001, by 2-way ANOVA (Sidak’s multiple comparisons test). (C) Recovery of infectious HCMV from DCs over time. DCs were pre-treated with A3A-targeting or negative control siRNA, and infected 48 h later by co-culture with HCMV-UL36-GFP-infected HFFF-His cells. DCs were purified using MACS and co-cultured with uninfected HFFFs after 72 h in triplicate. An Incucyte® was used to measure the total integrated GFP intensity (GCU x µ^2^/well) over the time course, normalised to timepoint 0. *p < 0.0101, ***p < 0.0001, ****p < 0.0001, by 2-way ANOVA (Sidak’s multiple comparisons test).

### Multiplexed proteomic comparison of plasma membrane protein changes in HCMV-infected and bystander primary DCs

Two biological replicates were performed for PM proteomic analysis: PM1 and PM2. As material was limiting, experiments were performed at the 72h timepoint only, facilitating analysis in a single multiplexed TMT experiment. Overall, 849 human proteins were quantified that had a Gene Ontology annotation indicative of a plasma membrane location (**Figure S4A, Methods**). Strong correlation between MACS- and FACS-enriched samples was again observed (**Figure S4B**). Uninfected, bystander and LPS-treated samples clustered separately from infected samples in both replicates, and proteins from bystander cells again changed in a similar manner to LPS-stimulated DCs (**Figure 6A, S4C**). Overall, 137 proteins were downregulated >2-fold and 153 proteins upregulated >2-fold compared to bystander cells (**Figures 6B-C, S4A, Table S6).** All data are shown in **Table S2**.

**Figure 6.**
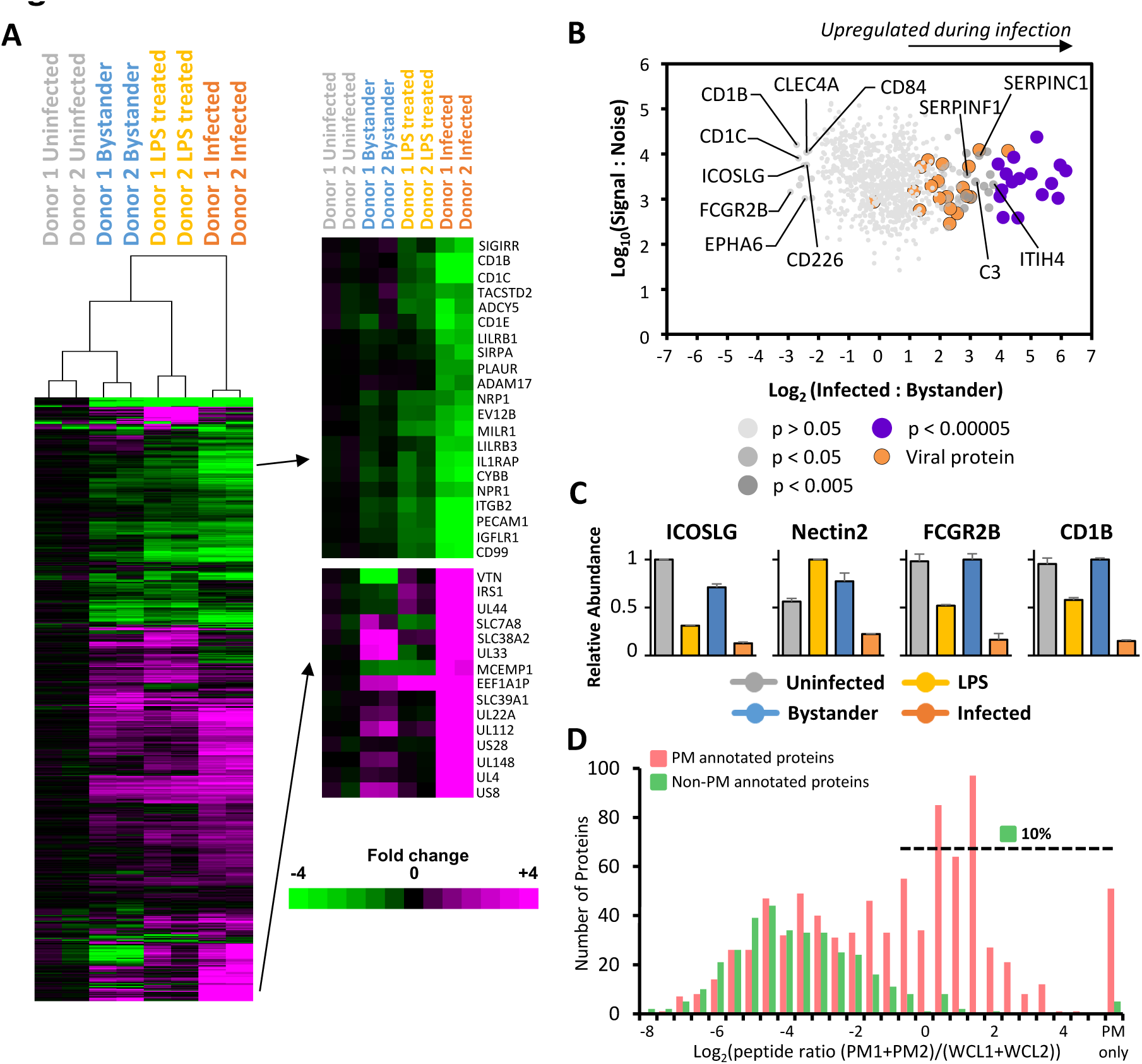
Proteomic analysis of plasma membrane proteins during HCMV infection in DCs. (A) Hierarchical cluster analysis of all 849 human proteins that had a Gene Ontology annotation indicative of a plasma membrane location, and all viral proteins from experiments PM1 or PM2. The fold change is shown compared to the average S:N of the two uninfected samples. An enlargement of two subclusters is shown on the right, including multiple proteins that were substantially up- or down-regulated during infection. (B) Scatterplot of all proteins from (A). The fold change and total S:N was averaged for proteins quantified in both experiments. Of note, this figure shows data in comparison to bystander samples, whereas (A) shows data in comparison to uninfected samples. Benjamini-Hochberg adjusted Significance A values were used to estimate p-values (see Methods). (C) Example protein profiles for downregulated PM proteins. (D) Filtering strategy to identify viral PM proteins. The plot shows the ratio of peptides (experiments PM1+PM2)/(experiments WCL1+WCL2) for every GO-annotated human protein quantified in experiment PM1 or PM2. 90% of proteins that were GO-defined non-PM had a ratio of <0.5; 93% of human proteins scoring above 0.5 were annotated as PM proteins, demonstrating the predictive value of this metric. Applying this filter, nine high confidence viral PM proteins were defined (**Figure S4E**).

Application of DAVID software to downregulated proteins suggested that a focus of HCMV at the DC plasma membrane is modulation of proteins important in adhesion, antigen presentation and antiviral immunity^22^. Multiple proteins with immunoglobulin-like domains, lectins and MHC class I molecules were downregulated. Interestingly, four of five CD1 family members were downregulated, which are structurally related to MHC proteins, have roles in presentation of lipid and glycolipid antigens but were not previously quantified in HFFFs^26^. Amongst downregulated molecules with other immune roles were Fc Gamma Receptor and Transporter (FCGRT), four high- or low-affinity Fc receptors (FCGR1A, 2B, 2C, 3A) and T-cell co-stimulators Nectin-2 and ICOSLG (**Figures 6C, S4D, Table S6A**). Proteins upregulated >2-fold were enriched in secreted proteins and endopeptidase inhibitors (**Figures 6C, S4D, Table S6B**).

Viral proteins present at the surface of an infected cell may be therapeutic antibody targets^27^. We detected a total of 19 viral proteins in experiments PM1 and PM2. Although our plasma membrane protein enrichment strategy provides a very substantial enrichment for PM glycoproteins, these preparations can be contaminated at a low level by abundant intracellular proteins such as certain viral proteins. We therefore employed a filtering strategy based on the ratio of PM and WCL peptides^18^, identifying nine candidate viral PM proteins with high confidence (**Figures 6D, S4E**). Of these, six corresponded to predictions from our previous analysis in HFFFs; one (UL148) was predicted to be present at the cell surface with high confidence in DCs, in comparison to a low confidence prediction in HFFFs, and two were not quantified at the HFFF cell surface (US28, UL4). US28 is a known surface therapeutic target^28^, and we recently identified UL4 as a secreted glycoprotein that inhibits TRAIL-mediated apoptosis and NK cell activation^29^.

### Viral manipulation of specific cell-surface proteins limits DC function

DCs play a pivotal role in the induction of adaptive immunity, with cell surface proteins functioning as key molecules governing interactions with other immune cells, such as CD4^+^ T-cells. The extensive virally-induced modulation of molecules involved in adhesion, antigen presentation and antiviral immunity at the DC plasma membrane suggested that HCMV-infected DCs might stimulate other immune cells inefficiently. To quantify this effect, we incubated HCMV-infected DCs with naïve CD4 T-cells that had been primed by an anti-CD3 antibody. T-cell proliferation was measured using a cell proliferation dye (CellTrace Violet) dilution assay, as a marker of activation. In this system, CD4^+^ T-cell proliferation was dependent on both stimulation via CD3 and the presence of DCs (**Figure 7A**).

**Figure 7.**
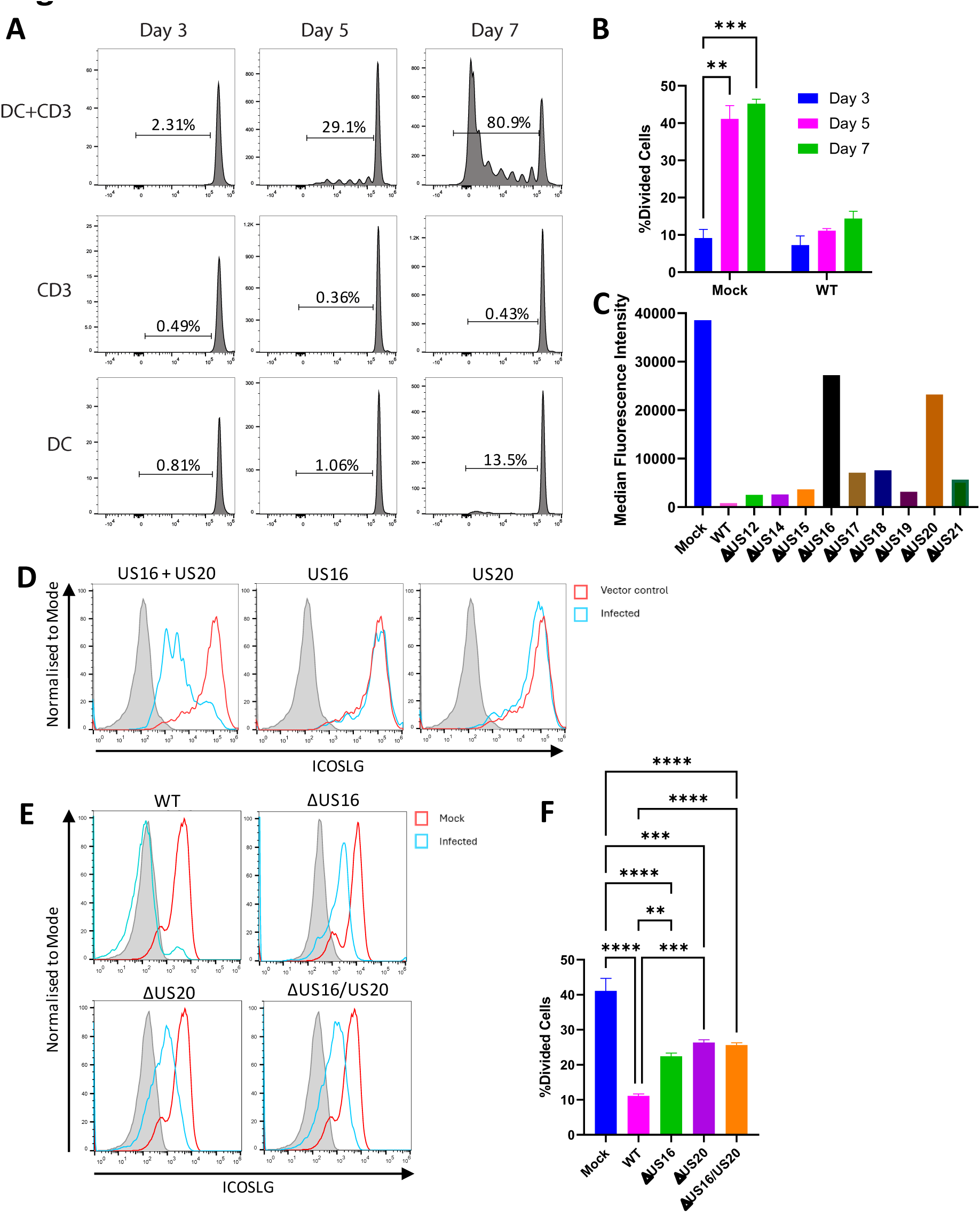
US16 and US20 inhibit cell-surface ICOSLG expression on DCs and limit CD4^+^ T-cell stimulation. (A) Development of an assay to detect DC stimulation of CD4^+^ T-cells. CD4^+^ cells were either unstimulated or stimulated with anti-CD3 antibody, then stained with CellTrace Violet and incubated alone or with DCs. At the indicated time points, samples were stained with an anti-CD4 antibody and analysed by flow cytometry. (B) HCMV limits DC-mediated CD4^+^ stimulation. CD4^+^ T-cells were stimulated with anti-CD3 antibody, then stained with CellTrace Violet and co-incubated with DCs that had previously been incubated with either mock- or HCMV-rCD2-infected HFFF-His cells, then purified by MACS. At the indicated time points, cells were stained with an anti-CD4 antibody and analysed by flow cytometry. (C) HCMV genes US16 and US20 reduce surface expression of exogenously-expressed ICOSLG. HF-CARs overexpressing ICOSLG (HF-ICOSLGs) were infected with HCMV or HCMV lacking genes from the US12 family at MOI = 5. After 72 h infection, cells were stained with anti-ICOSLG antibody and analysed by flow cytometry. (D) US16 and US20 co-operate to modulate ICOSLG. HF-ICOSLG were infected with adenoviruses expressing US16 or US20 at MOI 40. 72 h later, cells were stained with anti-ICOSLG antibody and analysed by flow cytometry. (E) US16 and US20 modulate endogenous ICOSLG expression in DCs. DCs were infected by co-culture with HCMV-rCD2 infected HFFF-His cells. Viruses used lacked US16, US20, or both. 24 h later newly infected DCs were purified using MACS, and at 48 h post-infection were stained with anti-ICOSLG antibody and analysed by flow cytometry. (F) Loss of US16 and/or US20 reduces the ability of HCMV to limit DC-mediated CD4^+^ T-cell proliferation. CD4^+^ T-cells were stimulated with anti-CD3 antibody, then stained with CellTrace Violet and co-incubated with DCs that had previously been incubated with either mock or HCMV-rCD2 infected HFFF-His cells, then purified by MACS. Viruses used were either HCMV-rCD2, or the same virus but lacking US16, US20, or both. At the indicated timepoints cells were stained with an anti-CD4 antibody and analysed by flow cytometry.

Mock-infected DCs stimulated extensive CD4^+^ T-cell proliferation, however this was almost completely abrogated upon incubation with purified HCMV-infected DCs, indicating that HCMV inhibits DC stimulation of adaptive immunity through direct modulation of the infected cell (**Figure 7B**). Amongst proteins downregulated at the plasma membrane by HCMV was ICOSLG, the ligand for the costimulatory molecule Inducible T-cell costimulatory (ICOS) (**Figures 6A-B**). ICOS is a member of the B7 family of co-stimulatory or co-inhibitory ligands and receptors. We have previously shown that another B7 family member, B7-H6 (a ligand for the activating receptor NKp30 on NK cells), is co-operatively targeted for lysosomal degradation by HCMV US18 and US20 in fibroblasts^30^. US18 and US20 are members of the US12 family, comprised of the US12 to US21 genes. We therefore determined whether ICOSLG might be targeted by proteins encoded in the same genetic region. We previously generated viral mutants lacking each individual member of the US12 family, however these can only infect fibroblasts due to genetic ablation of the pentameric complex. Since ICOSLG is not detectably expressed in fibroblasts, we overexpressed ICOSLG to identify the HCMV genes responsible for its loss. Knockout of either US16 or US20 rescued exogenously expressed ICOSLG in fibroblasts (**Figure 7C**). To determine whether these genes alone were sufficient for targeting of ICOSLG, HFFF overexpressing ICOSLG were transduced with Ad vectors expressing US16, US20, or both. Expression of either gene in isolation had only minimal effects, however significant downregulation was observed when both were expressed together (**Figure 7D**). Thus, US16 and US20 co-operate to target ICOSLG for removal from the cell surface, and are sufficient to carry out this activity in the absence of other viral genes. This effect was recapitulated in primary DCs expressing endogenous ICOSLG, where viruses lacking either US16 or US20 showed recovery of ICOSLG levels (**Figure 7E**). Importantly, DCs infected with HCMV lacking either US16, US20, or both genes, demonstrated an increased ability to stimulate CD4^+^ T-cells (**Figure 7F**), demonstrating that viral manipulation of specific host proteins on the surface of infected DCs directly inhibits the ability of these cells to prime adaptive immunity.

## Discussion

Herpesviruses achieve lifelong persistence in infected individuals by utilising a wide range of strategies to modulate innate and adaptive immunity. However, most studies of HCMV immune evasion have focused on a single cell type, fibroblasts. Although critical for viral replication during acute HCMV infection^31^, other cell types harbour a very different repertoire of functions and are equally important *in vivo*. In particular, monocyte lineage cells harbour latent virus, trigger lytic transcription upon differentiation into macrophages/DCs^32^, and facilitate haematogenous viral spread. Previous studies have used passaged HCMV strains and mixed populations of infected cells and bystanders to demonstrate that HCMV infection can impact a small number of individual DC proteins (e.g. MHC-II) and inhibit DC function^33–39^. However, because of this experimental design, it had been unclear to what extent these effects were due to direct modulation of the infected cell as opposed to influences of soluble mediators on bystander cells, and the complete repertoire of host functions that were modulated was unknown.

In this study we provide a comprehensive resource describing temporal changes in >120 viral and >8700 host proteins from HCMV-infected *ex-vivo* DCs, both within the cell and at the plasma membrane. In addition to highlighting host factors of particular importance in DCs based on their regulation by HCMV, our data and searchable database provide a comparative analysis to host protein regulation in fibroblasts. Unexpectedly, the correlation between global proteomic changes in both cell types was modest. 48% of proteins downregulated >2-fold in fibroblasts at 72 h were also downregulated >1.25-fold in DCs, and 69% of proteins downregulated >2-fold in DCs were also downregulated >1.25-fold in fibroblasts. A notable example of the differences between cell types included collagen proteins, which were globally downregulated in fibroblasts by HCMV and by other viruses^18,40,41^, yet minimally changed or upregulated in HCMV-infected DCs compared to bystanders. However, the group of proteins we previously identified as targeted for proteasomal or lysosomal degradation in HFFFs using three orthogonal proteomic/transcriptomic screens^21^ behaved much more similarly in both cell types. These included 35 proteins targeted with high confidence^21^ (HFFF:DC r^2^=0.47) and 133 proteins targeted with medium confidence^21^ (HFFF:DC r^2^ = 0.23). Re-examining our original data to identify a third group of 402 proteins degraded with lower confidence, these still correlated better between HFFFs and DCs (r^2^=0.18) than the general correlation between all protein changes (r^2^=0.10). This suggests that additional factors on the ‘low confidence’ list are also degraded by HCMV, expanding the potential list of HCMV’s targets. Interestingly, this also suggests that another measure in the confidence a given protein is degraded can be derived from similar regulation in diverse cell types, and that selection of future ‘hits’ for detailed characterisation might be chosen on the same basis. Importantly however, the repertoire of host proteins expressed by fibroblasts differed substantially from DCs, and as a result, there were multiple aspects of the virus-host interaction that were only revealed in individual cell types.

Using passaged cell-free virus, studies differ as to the efficiency of DC infection, with earlier reports showing very high levels of infection^42,43^, while more recent studies found that (unlike in HFFFs), only a proportion of DCs become infected^44^. This was attributed to stochastic heterogeneity in the virus:host interaction. Specifically, some cells mount a more rapid innate immune response and prevent infection, while others express viral evasins early enough to counteract this response enabling infection to proceed^44^. Virus strain and the route of infection can substantially alter the outcome of infection in DCs; we previously found that cell-cell spread of WT-HCMV into DCs enables the virus to overcome innate immunity more rapidly following pre-treatment with interferon as compared with cell-free infection^45^. We also showed that WT-HCMV encodes higher levels of the pentameric glycoprotein complex than passaged strains, which inhibits virus spread in the absence of antibody, but enables spread to proceed in the presence of neutralising antibodies^14,45^. Thus, multiple conditions impact DC infection, with some phenotypes only being revealed in certain settings. It is likely that infection by the cell-cell route occurs in a more inflammatory milieu then cell-free infection, due to release of interferons from the infected fibroblasts (supported by our data on bystander cell protein changes). Nevertheless, efficiency of cell-cell infection by wildtype virus in the present analysis was similar to the more recent studies in the literature using cell-free virus, with 20-40% of DCs becoming infected^45^.

We now show that the kinetics of viral gene expression following cell-cell transfer are also delayed compared to HFFFs. Using the temporal profiles from our previous analysis of infected fibroblasts^18^, the Tp4 class of proteins was delayed by 24 h, and Tp5 proteins were delayed by 48 h, with a concomitant delay in VAC formation. DCs thus encode restriction mechanisms that are absent from HFFF and operate at both the early^44^ and late phases of the lifecycle, with late-phase inhibition mediated at least in part by A3A, one of a family of cytidine deaminases that have recently been shown to have inhibitory effects on a range of RNA and DNA viruses^46^ following interferon induction. A recent study showed that HCMV infection in decidual tissue can lead to upregulation of A3A as a result of interferon release from infected cells^47^, which we also saw in bystander DCs, but not in HFFFs^18^. HCMV has previously been shown to be sensitive to A3A following exogenous overexpression in epithelial cells, where inhibition occurred for all temporal classes of HCMV genes^47^. Our data now demonstrates that endogenous A3A forms a natural antiviral restriction factor for HCMV in DCs, and is capable of delaying (but not completely preventing) the cascade of viral gene expression from Tp4 onwards following cell-cell transfer of a wild-type virus. Another study examining overexpressed APOBEC family members in epithelial cells concluded that HCMV is capable of relocalising A3B but not A3A, although whether this relocalisation affected function was not tested^48^. Our proteomic data now suggest that HCMV encodes modulators of both A3A and A3B since expression in infected cells was dramatically limited compared to bystanders. Monocyte-derived DCs are not the only cell type to restrict HCMV lytic infection. Following cell-free infection, plasmacytoid and CD11c^+^ DCs, and immature Langerhans DCs^49,50^ are resistant to productive infection, and trophoblast stem cells, extravillous trophoblasts, and syncytiotrophoblasts have a block just after the IE stage of infection^51^. This may also be explained by differential expression of restriction factors compared to fibroblasts. In addition to the poor efficiency of lytic infection in monocyte-derived DCs that we show here, reactivation of latent infection during DC differentiation can also be inefficient^32^. It will be interesting to determine whether A3A similarly restricts reactivation from latency, especially given that reactivation occurs in response to inflammatory stimuli, which our data suggests can upregulate A3A^52^.

Previous studies of viral immune evasins have not only influenced our understanding of pathogen immune evasion, but have revealed fundamental mechanisms of immune system function such as the requirements for immune-synapse formation^53^, and novel aspects of peptide presentation^54^. HCMV has the largest genome of any human virus, encoding over 170 ORFs, of which only 41-45 are essential for replication^55,56^, with the remainder encoding accessory functions. Although the function of many of these are unknown, over 19 genes and one microRNA target cell-surface ligands for NK cells^3,29,57^, and at least four genes target cell-surface MHC-I to limit CD8^+^ T-cell activation^7^. Our data now reveal that in addition to genes targeting MHC-II^58^, the virus is also likely to encode a wide range of additional modulators that target proteins that are absent from HFFFs but present in APCs, and that are capable of altering APC function. One of these mechanisms operates via US16 and US20, which downregulate ICOSLG on the surface of infected cells, and further inhibit the ability of DCs to stimulate CD4^+^T cells even despite active viral modulation of MHC-II. However, given the number of additional proteins altered on the infected cell surface, it is likely that there are many more DC-specific immune evasins. HCMV is unique amongst viruses in the extent to which it imprints on lifelong host immunity^6^, including adaptive immunity, and many of the modulations of the DC surface could underpin these observations.

In addition to modulation of host immunity, our analysis also revealed the existence of viral antigens on the surface of infected cells. We have previously shown that infection of HFFF leads to cell-surface expression of viral proteins that are potent targets for antibody-dependent cellular cytotoxicity (ADCC)^27^. Interestingly, viral immune evasins expressed early in infection, including UL141 and UL16, were superior targets compared to the structural proteins expressed late in infection^27,59,60^ such as gB, which are commonly used in subunit vaccines. Our DC data now demonstrates that these potent ADCC targets are also present on DC surfaces from early times. Given the delayed nature of the replication cycle in DCs, it may be even more critical to target these early-expressed cell-surface antigens rather than late-expressed structural antigens, to provide the longest possible window for antibody-dependent cellular immunity to respond to infection of APCs and control infection.

In summary, we have developed novel technologies that permit routine isolation of purified APCs following infection with wild-type HCMV by a clinically relevant route. Proteomic analysis revealed a wide range of previously unknown protein changes in infected cells that may dictate the outcome of infection, many involving factors that are not expressed in cell types commonly used for modelling HCMV pathogenesis. These proteins included host restriction factors capable of preventing or delaying the viral lifecycle, and cell surface proteins capable of influencing the induction of protective immunity.

## Methods & Materials

### Reagents and Tools

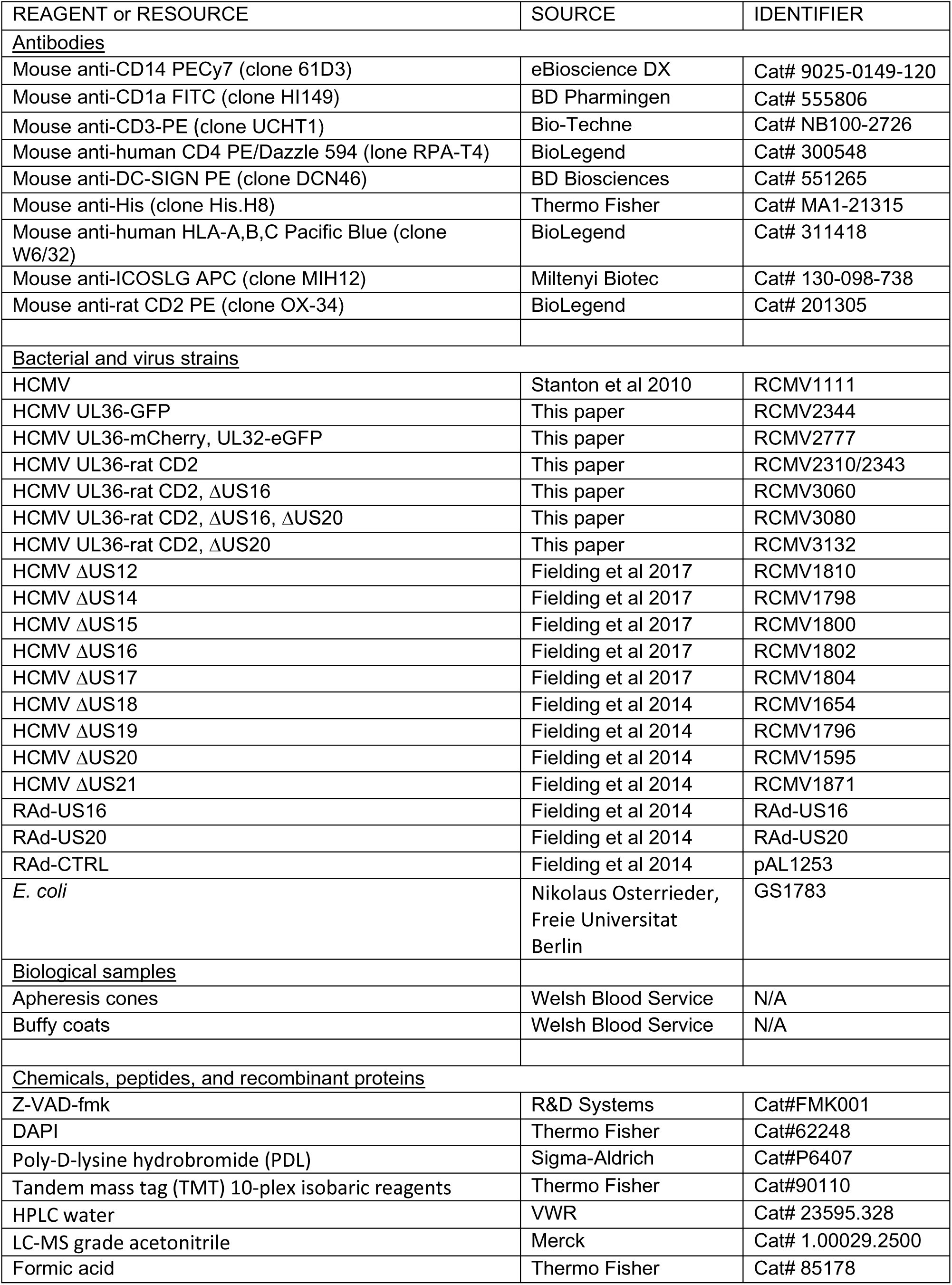
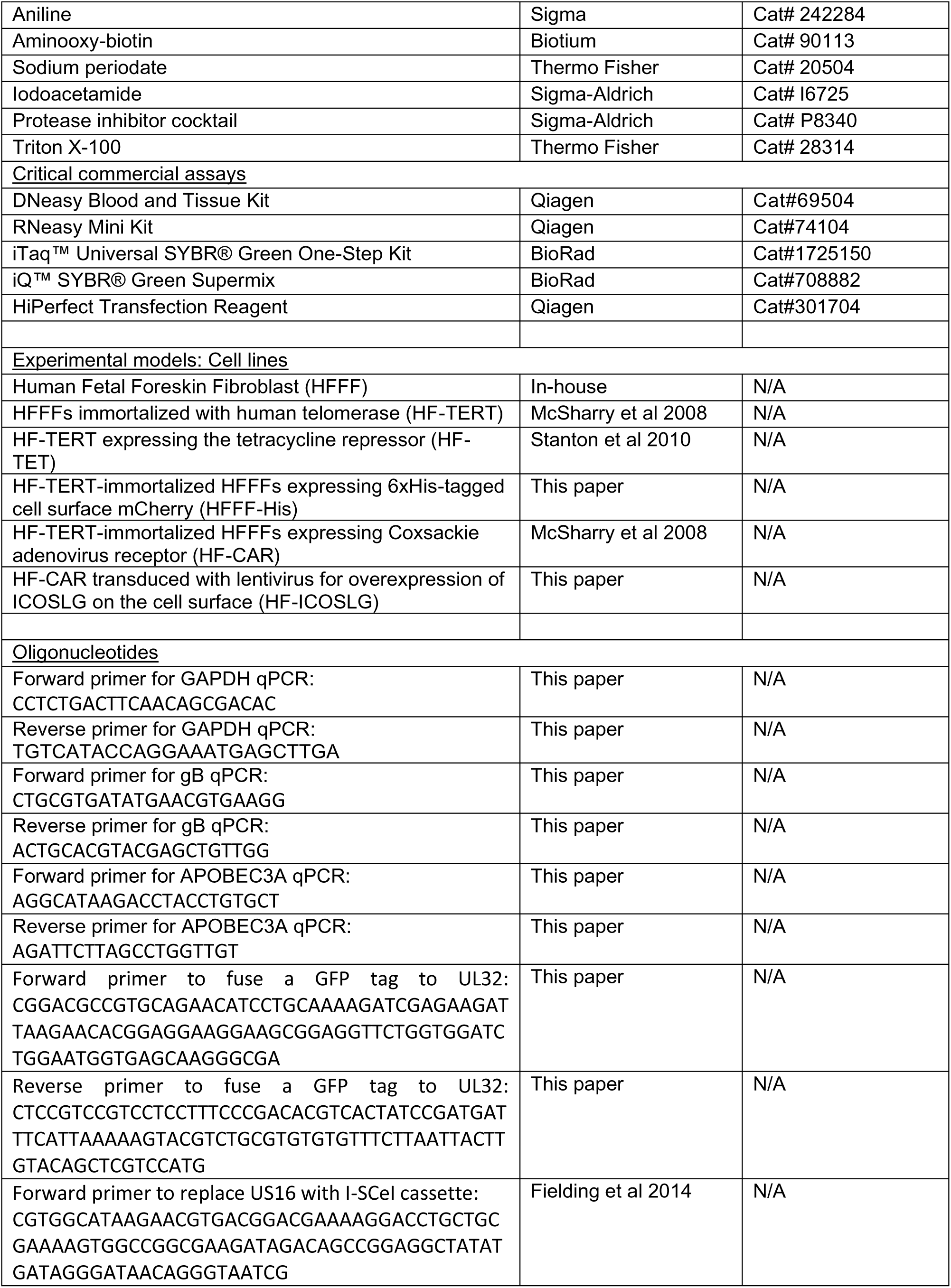
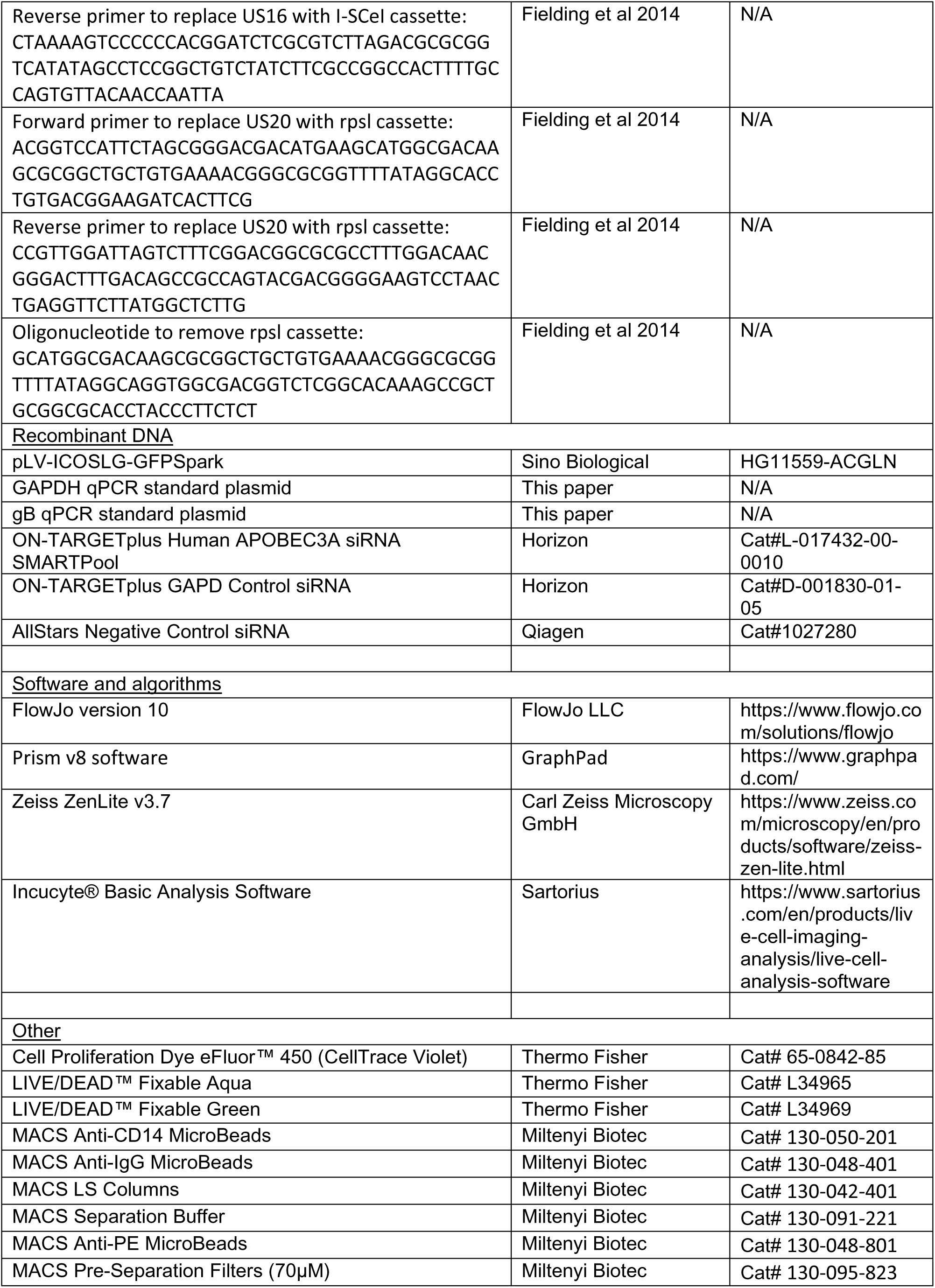

#### Cells and viruses

Human foetal foreskin fibroblasts (HFFFs) immortalised with human telomerase reverse transcriptase (HF-TERTs)^61,62^, and HF-TERTs expressing the tetracycline repressor (HF-TETs)^17^, have been described before. HF-TERTs expressing 6xHis-tagged cell-surface mCherry (HFFF-His) were generated by gene synthesizing a codon optimised ORF encoding mCherry with a N-terminal signal sequence followed by a 6xHis tag, and a GPI anchor derived from CD55 at its C-terminus. This ORF was cloned into the retroviral vector pMXS-IP^63^ using XhoI/NotI restriction sites, then retrovirus produced by transfecting 293 Phoenix cells. HF-TERTs were then infected with the retrovirus using retronectin, and selected using puromycin (1mg/ml). HF-TERT-immortalized HFFFs expressing Coxsackie adenovirus receptor (HF-CAR)^61^ were transduced with a lentivirus vector expressing ICOSLG-GFP for the purpose of overexpressing ICOSLG on a fibroblast line. All cells were maintained under standard conditions in DMEM (Sigma-Aldrich) supplemented with 10% FCS.

All HCMV viruses were derived from bacterial artificial chromosomes (BACs) containing the complete Merlin HCMV genome^17,64^. Viruses containing a GFP tag linked to UL36 by a P2A peptide have been described before^21^. To insert a mCherry or truncated RatCD2 sequence in place of the GFP tag, codon optimised ORFs were gene synthesized, and cloned in to the Merlin BAC by en-passant mutagenesis^65^, replacing the original GFP ORF. To fuse a GFP tag to UL32, the eGFP ORF was PCR amplified using primers incorporating a linker sequence (GSGGSGGSG; primers CGGACGCCGTGCAGAACATCCTGCAAAAGATCGAGAAGATTAAGAACACGGAGGAAGGAAGCGGAGGTTCTGGTGGATCTGGAATGGTGAGCAAGGGCGA and CTCCGTCCGTCCTCCTTTCCCGACACGTCACTATCCGATGATTTCATTAAAAAGTACGTCTGCGTGTGTGTTTCTTAATTACTTGTACAGCTCGTCCATG), and fused to the end of the UL32 ORF by en passant mutagenesis. Deletion of US16 was carried out by en-passant mutagenesis^65^. Primers used were CGTGGCATAAGAACGTGACGGACGAAAAGGACCTGCTGCGAAAAGTGGCCGGCGAAGATAGACAGCCGGAGGCTATATGATAGGGATAACAGGGTAATCG and CTAAAAGTCCCCCCACGGATCTCGCGTCTTAGACGCGCGGTCATATAGCCTCCGGCTGTCTATCTTCGCCGGCCACTTTTGCCAGTGTTACAACCAATTA. Deletion of US20 was carried out by recombineering using a rpsl cassette^66^; primers used to insert the cassette were ACGGTCCATTCTAGCGGGACGACATGAAGCATGGCGACAAGCGCGGCTGCTGTGAAAACGGGCGCGGTTTTATAGGCACCTGTGACGGAAGATCACTTCG, CCGTTGGATTAGTCTTTCGGACGGCGCGCCTTTGGACAACGGGACTTTGACAGCCGCCAGTACGACGGGGAAGTCCTAACTGAGGTTCTTATGGCTCTTG, while the following primer was used to delete the cassette along with US20: GCATGGCGACAAGCGCGGCTGCTGTGAAAACGGGCGCGGTTTTATAGGCAGGTGGCGACGGTCTCGGCACA AAGCCGCTGCGGCGCACCTACCCTTCTCT. Viruses were generated by transfection of the BAC into HF-TETs, then virus harvested from the supernatant, pelleted to remove cellular debris and concentrated by centrifugation, before being titrated on primary HFFFs. Adenovirus vectors were constructed as previously described^66^. All viruses underwent whole genome sequencing following reconstitution to ensure the expected modification was present and that no second-site mutations had occurred^15^. A list of viruses used in the publication can be found in **Reagents and Tools Table**.

Peripheral blood mononuclear cells (PBMC) were obtained from healthy volunteers, or from apheresis cones, and isolated from whole blood by Histopaque centrifugation. CD14^+^ monocytes were isolated by MACS using the human CD14^+^ isolation kit (Miltenyi Biotec). CD14^+^ monocytes were differentiated into DCs over 6 days by culturing in RPMI (Sigma-Aldrich) supplemented with 10% FCS, 100ng/ul IL-4 (Peprotech), 100ng/ul GM-CSF (Peprotech) and 50nM β-mercaptoethanol (Gibco), unless they were to be used for siRNA knockdown in which case they were cultured in IMDM (Thermo Fisher). Dendritic cells were phenotyped by flow cytometry using antibodies against DC-SIGN, CD1a and CD14 (**Reagents and Tools Table**). To transfer virus from infected HFFF-His cells into DCs, media was removed from the HFFF-His cells and replaced with 10% RPMI (or IMDM) containing the DCs for 24h.

#### Separation of subpopulations following co-culture

Following 24h of co-culture, supernatant was collected, then monolayers were washed, and detached cells combined with the supernatant. Detached and suspension cells were stained with anti-His antibody, followed by anti-mouse-IgG magnetic beads. Labelled and unlabelled cells were then separated by magnetic activated cell-sorting (MACS; Miltenyi Biotec). Unlabelled cells (DCs) were then rested for 2h before being stained with anti-rat CD2-PE antibody, followed by anti-PE magnetic beads. Labelled and unlabelled cells were then again separated by MACS (Miltenyi Biotec).

#### Flow cytometry

Cells were washed in PBS and stained with primary antibody (**Reagents and Tools Table**) for 15 min at 4°C; if the antibody was conjugated to a fluorophore then cells were washed in PBS after incubation before being fixed with 4% paraformaldehyde (PFA). Unconjugated antibodies were washed and incubated with the secondary antibody for 15 min at 4°C before being washed and fixed. All data was acquired using an Attune Nxt Flow Cytometer (Thermo Fisher) unless otherwise stated, and analysed using FlowJo Software version 10 (FlowJo LLC).

#### qPCR & RT-qPCR

To ensure dying cells did not skew results, the pan caspase inhibitor Z-VAD was added to cultures. DNA was isolated from cells using the DNeasy Blood & Tissue Kit (Qiagen 69504) and stored at -20°C, for dendritic cells DNA extraction was performed following centrifugation over Histopaque to remove dead/dying cells from the population.

The standards used for qPCR experiments were a series of 10-Fold dilutions of DNA from plasmids containing gB and GAPDH. Reaction mixes for qPCR each comprised 20μl - 10μl SYBR Green (BioRad), 0.4μl each of the forward and reverse primers (**Reagents and Tools Table**), and 100ng of sample DNA diluted in DNAse and RNAse-free water, adjusted to a final volume of 20μl with DNAse and RNAse-free water. Each sample was added to an Applied Biosystems™ MicroAmp™ EnduraPlate™ Optical 96-Well Clear Reaction Plate (Thermo Fisher) in triplicate for each reaction, the plate was sealed, vortexed and centrifuged briefly before being placed into a QuantStudio 3 Real-Time qPCR System machine. The thermocycling programme, Comparative C_T_ with Melt, is presented in **Table S9**. Once the programme was complete, the data was analysed using Thermofisher Standard Curve or Relative Quantitation software for qPCR and RT-qPCR, respectively.

For RT-qPCR, RNA was isolated from cells using the RNeasy Plus Mini Kit (Qiagen), samples were stored at -80°C. iTaq™ Universal SYBR® Green One-Step Kit (Bio-Rad) was used for RT-qPCR. The 20μl reaction mix was set up using the RT-qPCR primers found in **Reagents and Tools Table**, according to the manufacturer’s protocol. Reactions were set up in triplicate in a 96-well plate and placed into a thermocycler as above.

#### Knock-down of cellular proteins using siRNA

To knockdown cellular proteins in primary DCs, the cells underwent one round of transfection being harvested for validation by RT-PCR two days following transfection. The reaction mix consisted of 2μl HiPerfect transfection reagent (Qiagen), 150nM siRNA (**Reagents and Tools Table**) and 45μl 0% IMDM, which was incubated for 15 min in a 96-well plate. After incubation, 100,000 DCs per 150μl 10% IMDM were added to the reaction mix for reverse transfection. Volumes were scaled up accordingly when required. RT-qPCR was used to measure the fold change of A3A compared to GAPDH (housekeeping gene). The ΔΔCT CT method was used to calculate the expression change following treatment of DCs with A3A siRNA (**Table S8**).

#### Immunofluorescence

DCs (infected, bystanders and uninfected) were seeded into a glass-bottom 24-well plate which had been pre-treated with poly-D-lysine hydrobromide (PDL, Sigma-Aldrich) for 1-24 h. DCs were fixed with 4% PFA for 15 min, washed, and permeabilised with 0.5% NP-40 for a further 15 min before being washed and stained with DAPI (300nM, Invitrogen). DABCO mounting media was added before being visualised using the Zeiss LSM800 confocal laser scanning microscope with a 63X oil lens.

#### Incucyte® live-cell analysis system

HF-TERTs were seeded into a 24-well plate and co-cultured with A3A siRNA treated infected DCs (HCMV UL36-GFP). Co-cultures were visualised using the Incucyte® SX5, the whole well was scanned with a 4X lens, with both brightfield and green image channels. Wells were scanned every 12 h for 9 days post co-culture. Incucyte® Basic Analysis Software was used to measure the Total Integrated Intensity (GCU x µ^2^/well) over the time course. Total Integrated Intensity was normalised to timepoint 0 for each condition.

#### Assessment of DC-mediated CD4^+^ T-cell proliferation

Naïve CD4^+^ T-cells were purified from healthy donor PBMCs by MACS (Miltenyi Biotec), then stimulated for 24h with anti-CD3 antibody (clone UCHT1, 25ng/ml). Cells were stained with CellTrace Violet (20 nM), then incubated with DCs previously incubated for 24 h with HCMV-rCD2, or Mock-infected, HFFF-His, then purified by MACS. Cells were harvested three, five, or seven days later, stained with anti-CD4 PE/Dazzle, and analysed by flow cytometry.

#### Enrichment of DCs for proteomic analysis

After separation of HFFF-His cells and DCs, and then of infected and bystander DCs, cells were pelleted by centrifugation at 500 g for 5 min. The pellet was resuspended in 1ml PBS and layered over 3ml Histopaque in a 15 ml Falcon then centrifuged at 2000 rpm for 20 min with the brake off. DCs were then removed from the interface, washed in PBS, and pelleted again.

#### Plasma membrane protein enrichment for proteomics

Biotinylation of the DCs was performed using a two-step method where oxidation and aminooxy-biotinylation are carried out separately^67^. The DC pellet was resuspended in 1mM sodium periodate (Thermo Fisher) and incubated on a Falcon Roller at 4°C for 20 min in the dark. The reaction was quenched with glycerol to 1 mM final concentration, then cells were pelleted and washed twice in ice-cold PBS pH 7.4. A biotinylation mix was generated within the 10 minutes prior to addition to the DCs. This consisted of 100 mM aminooxy-biotin (Biotium), 10 mM aniline (Sigma-Aldrich) and 5% v/v filtered foetal calf serum in ice-cold PBS pH 6.7. All PBS was aspirated from the DC pellet prior to biotinylation at 4°C on a Falcon Roller for a further 30 min. DCs were centrifuged, washed twice in ice-cold PBS pH 7.4 then lysed in lysis buffer (1% Triton X-100 (high purity, Thermo Fisher); 150 mM NaCl, 1× protease inhibitor (complete, without EDTA (Roche)), 5 mM iodoacetamide (Sigma-Aldrich), 10 mM Tris-HCl) on ice for 30 min. Samples were then centrifuged for 5 min at 13,000 g at 4°C. The supernatant was transferred to a fresh Eppendorf, and centrifuged a further two times, transferring supernatant to a fresh Eppendorf between each spin. Lysates were snap frozen in an ethanol/dry ice bath and stored at -80°C.

Biotinylated glycoproteins were enriched with high affinity streptavidin agarose beads (Thermo Fisher) and washed extensively using a vacuum manifold and Poly-Prep columns (BioRad). Washing was initially with lysis buffer, then 0.5% SDS and then urea. Captured protein was reduced with DTT, alkylated with iodoacetamide (IAA, Sigma-Aldrich) and digested on-bead with trypsin (Promega) in 100 mM HEPES pH 8.5 for 3h. Tryptic peptides were collected.

#### Whole cell lysate protein enrichment for proteomics

The DC pellet was washed twice with PBS, and 200 μl lysis buffer added (6M Guanidine/200 mM HEPES pH 8.5). Cells were vortexed extensively then sonicated. Cell debris was removed by centrifuging at 21,000 g for 10 min twice. For half of each sample, dithiothreitol (DTT) was added to a final concentration of 5 mM and samples were incubated for 20 mins. Cysteines were alkylated with 14 mM iodoacetamide and incubated 20 min at room temperature in the dark. Excess iodoacetamide was quenched with DTT for 15 mins. Samples were diluted with 200 mM HEPES pH 8.5 to 1.5 M Guanidine followed by digestion at room temperature for 3 h with LysC protease at a 1:100 protease-to-protein ratio. Samples were further diluted with 200 mM HEPES pH 8.5 to 0.5 M Guanidine. Trypsin was then added at a 1:100 protease-to-protein ratio followed by overnight incubation at 37°C. The reaction was quenched with 5% formic acid, then centrifuged at 21,000 g for 10 min to remove undigested protein. Peptides were subjected to C18 solid-phase extraction (SPE, Sep-Pak, Waters) and vacuum-centrifuged to near-dryness.

#### Peptide labelling with tandem mass tags (TMT)

In preparation for TMT labelling, desalted peptides were dissolved in 200 mM HEPES pH 8.5. For WCL samples, peptide concentration was measured by microBCA (Pierce), and 25 μg of peptide labelled with TMT reagent. 10-plex TMT reagents (0.8mg, Thermo Fisher) were dissolved in 43 µl of anhydrous acetonitrile, and 5 µl was added to the peptide samples at a final concentration of 30% acetonitrile (v/v). Following incubation for 1 h at room temperature, the reaction was quenched with hydroxylamine to a final concentration of 0.05% (v/v). Small aliquots of TMT-labelled samples were combined 1:1:1:1:1:1:1:1:1 (sample WCL1) or 1:1:1:1:1:1:1:1 (samples WCL2 and PM sample), vacuum centrifuged to near dryness and desalted using a StageTip^68^. An unfractionated singleshot was analysed initially to ensure similar peptide loading across each TMT channel, thus avoiding the need for excessive electronic normalization. If normalisation factors were >0.5 and <2, data for each singleshot experiment were analysed with data for the corresponding fractions to increase the overall number of peptides quantified. Otherwise, samples were re-mixed in appropriate proportions, a ‘singleshot’ experiment repeated and data for this experiment analysed with data for the corresponding fractions after confirming normalisation factors were >0.5 and <2. Normalisation is discussed in ‘Data Analysis’, and high pH reversed-phase (HpRP) and strong cation exchange (SCX) fractionation are discussed below. Details of individual sample labelling, and mass spectrometry analyses are described in **Table S7**.

#### Offline tip-based strong cation exchange SCX fractionation

A protocol for solid-phase extraction based SCX peptide fractionation was previously modified for small peptide amounts^18^. Briefly, 10 mg of PolySulfethyl A bulk material (Nest Group Inc) was loaded on to a fritted 200 µl tip in 100% acetonitrile using a vacuum manifold. The SCX material was conditioned slowly with 2x 400 µl SCX buffer A (7mM KH_2_PO_4_, pH 2.65, 30% Acetonitrile), 400 µl SCX buffer B (7mM KH_2_PO_4_, pH 2.65, 350mM KCl, 30% Acetonitrile) and then 4x 400 µl SCX buffer A. Dried peptides were resuspended in 400 µl SCX buffer A and added to the tip under vacuum, with a flow rate of ∼150 µl/min. The tip was then washed with 150 µl SCX buffer A. Fractions were eluted in 150 µl washes using SCX buffer at increasing K+ concentrations (flow-through, 10, 25, 40, 60, 90, 150mM KCl), vacuum-centrifuged to near dryness then desalted using StageTips^68^.

#### Offline HpRP fractionation

TMT-labelled tryptic peptides were subjected to HpRP fractionation using an Ultimate 3000 RSLC UHPLC system (Thermo Fisher) equipped with a 2.1 mm internal diameter x 25 cm long, 1.7 µm particle Kinetix Evo C18 column (Phenomenex). The mobile phase consisted of A: 3% acetonitrile (MeCN), B: MeCN and C: 200 mM ammonium formate pH 10. Isocratic conditions were 90% A / 10% C, and C was maintained at 10% throughout the gradient elution. Separations were conducted at 45°C. Samples were loaded at 200 µl/min for 5 min. The flow rate was then increased to 400 µl/min over 5 min, after which the gradient elution proceed as follows: 0-19% B over 10 min, 19-34% B over 14.25 min, 34-50% B over 8.75 min, followed by a 10 minwash at 90% B. UV absorbance was monitored at 280 nm and 15 s fractions were collected into 96 well microplates using the integrated fraction collector. Fractions were recombined orthogonally in a checkerboard fashion, combining alternate wells from each column of the plate into a single fraction, and commencing combination of adjacent fractions in alternating rows. Wells were excluded prior to the start or after the cessation of elution of peptide-rich fractions, as identified from the UV trace. This yielded two sets of 12 combined fractions, A and B, which were dried in a vacuum centrifuge and resuspended in 10 µl MS solvent (4% MeCN / 5% formic acid) prior to LC-MS3. 12 set ‘A’ fractions were used for MS analysis of all experiments.

#### LC-MS3

Mass spectrometry data were acquired using an Orbitrap Lumos. Loading solvent was 0.1% FA, analytical solvent A: 0.1% FA and B: 80% MeCN + 0.1% FA. All separations were carried out at 55°C. Samples were loaded at 5 µL/min for 5 min in loading solvent before beginning the analytical gradient. The following gradient was used: 3-7% B over 3 min, 7-37% B over 173 min, followed by a 4 min wash at 95% B and equilibration at 3% B for 15 min. Each analysis used a MultiNotch MS3-based TMT method^21,69^. The following settings were used: MS1: 380-1500 Th, 120,000 Resolution, 2x10^5^ automatic gain control (AGC) target, 50 ms maximum injection time. MS2: Quadrupole isolation at an isolation width of m/z 0.7, CID fragmentation (normalised collision energy (NCE) 35) with ion trap scanning in turbo mode from m/z 120, 1.5x10^4^ AGC target, 120 ms maximum injection time. MS3: In Synchronous Precursor Selection mode the top 6 MS2 ions were selected for HCD fragmentation (NCE 65) and scanned in the Orbitrap at 60,000 resolution with an AGC target of 1x10^5^ and a maximum accumulation time of 150 ms. Ions were not accumulated for all parallelisable time. The entire MS/MS/MS cycle had a target time of 3 s. Dynamic exclusion was set to +/- 10 ppm for 70 s. MS2 fragmentation was trigged on precursors 5x10^3^ counts and above.

### Quantification and Statistical Analysis

#### Data analysis

Mass spectra were processed using a Sequest-based software pipeline for quantitative proteomics, “MassPike”, through a collaborative arrangement with Professor Steven Gygi’s laboratory at Harvard Medical School. MS spectra were converted to mzXML using an extractor built upon Thermo Fisher’s RAW File Reader library (version 4.0.26). In this extractor, the standard mzxml format has been augmented with additional custom fields that are specific to ion trap and Orbitrap mass spectrometry and essential for TMT quantitation. These additional fields include ion injection times for each scan, Fourier Transform-derived baseline and noise values calculated for every Orbitrap scan, isolation widths for each scan type, scan event numbers, and elapsed scan times. This software is a component of the MassPike software platform and is licensed by Harvard Medical School.

A combined database was constructed from (a) the human Uniprot database (26^th^ January, 2017), (b) the HCMV strain Merlin Uniprot database, (c) all additional non-canonical human cytomegalovirus ORFs described by Stern-Ginossar et al^70^, (d) a six-frame translation of HCMV strain Merlin filtered to include all potential ORFs of ≥8 amino acids (delimited by stop-stop rather than requiring ATG-stop) and (e) common contaminants such as porcine trypsin and endoproteinase LysC. ORFs from the six-frame translation (6FT-ORFs) were named as follows: 6FT_Frame_ORFnumber_length, where Frame is numbered 1-6, and length is the length in amino acids. The combined database was concatenated with a reverse database composed of all protein sequences in reversed order. Searches were performed using a 20 ppm precursor ion tolerance. Fragment ion tolerance was set to 1.0 Th. TMT tags on lysine residues and peptide N termini (229.162932 Da) and carbamidomethylation of cysteine residues (57.02146 Da) were set as static modifications, while oxidation of methionine residues (15.99492 Da) was set as a variable modification.

To control the fraction of erroneous protein identifications, a target-decoy strategy was employed^71^. Peptide spectral matches (PSMs) were filtered to an initial peptide-level false discovery rate (FDR) of 1% with subsequent filtering to attain a final protein-level FDR of 1%. PSM filtering was performed using a linear discriminant analysis, as described previously^71^. This distinguishes correct from incorrect peptide IDs in a manner analogous to the widely used Percolator algorithm (https://noble.gs.washington.edu/proj/percolator/), though employing a distinct machine learning algorithm. The following parameters were considered: XCorr, ΔCn, missed cleavages, peptide length, charge state, and precursor mass accuracy.

Protein assembly was guided by principles of parsimony to produce the smallest set of proteins necessary to account for all observed peptides (algorithm described in^71^. Where all PSMs from a given HCMV protein could be explained either by a canonical gene or non-canonical ORF, the canonical gene was picked in preference. In certain cases, PSMs assigned to a non-canonical or 6FT-ORF were a mixture of peptides from the canonical protein and the ORF. This most commonly occurred where the ORF was a 5’-terminal extension of the canonical protein (thus meaning that the smallest set of proteins necessary to account for all observed peptides included the ORFs alone). In these cases, the peptides corresponding to the canonical protein were separated from those unique to the ORF, generating two separate entries. In a single case, PSM were assigned to the 6FT-ORF 6FT_6_ORF1202_676aa, which is a 5’-terminal extension of the non-canonical ORF ORFL147C. The principles described above were used to separate these two ORFs.

Proteins were quantified by summing TMT reporter ion counts across all matching peptide-spectral matches using ”MassPike”, as described previously^69^. Briefly, a 0.003 Th window around the theoretical m/z of each reporter ion (126, 127n, 127c, 128n, 128c, 129n, 129c, 130n, 130c, 131n, 131c) was scanned for ions, and the maximum intensity nearest to the theoretical m/z was used. The primary determinant of quantitation quality is the number of TMT reporter ions detected in each MS3 spectrum, which is directly proportional to the signal-to-noise (S:N) ratio observed for each ion. Conservatively, every individual peptide used for quantitation was required to contribute sufficient TMT reporter ions so that each on its own could be expected to provide a representative picture of relative protein abundance^69^. An isolation specificity filter with a cutoff of 50% was additionally employed to minimise peptide co-isolation^69^. Peptide-spectral matches with poor quality MS3 spectra (more than 9 TMT channels missing and/or a combined S:N ratio of less than 100 across all TMT reporter ions) or no MS3 spectra at all were excluded from quantitation. Peptides meeting the stated criteria for reliable quantitation were then summed by parent protein, in effect weighting the contributions of individual peptides to the total protein signal based on their individual TMT reporter ion yields. Protein quantitation values were exported for further analysis in Excel.

For protein quantitation, reverse and contaminant proteins were removed, then each reporter ion channel was summed across all quantified proteins and normalised assuming equal protein loading across all channels (these normalised data are presented in **Table S2**). For further analysis and display in Figures, fractional TMT signals were used (i.e. reporting the fraction of maximal signal observed for each protein in each TMT channel, rather than the absolute normalized signal intensity). This effectively corrected for differences in the numbers of peptides observed per protein. For **Figure 6**, PM proteins were defined as by a GO annotation of “plasma membrane”, “cell surface”, “extracellular” or “short GO”^67^. All analyses for this Figure was performed on data filtered to include only those proteins with one of these GO annotations.

Hierarchical centroid clustering based on uncentered Pearson correlation, and k-means clustering were performed using Cluster 3.0 (Stanford University) and visualised using Java Treeview (http://jtreeview.sourceforge.net) unless otherwise noted.

#### IBAQ analysis

The intensity-based absolute quantification (IBAQ) method was adapted from the original description^72^ as we previously described for WT HCMV strain Merlin infection of HF-TERT cells at 24, 48 and 72 h post infection^25^. The maximum MS1 precursor intensity for each quantified viral peptide was determined for each experiment, and a summed MS1 precursor intensity for each viral protein across all matching peptides was calculated. To determine the proportion of the summed intensity that arose at 24, 48 and 72 h PI, the summed intensity was adjusted in proportion to normalized TMT values: (24 h + 48 h + 72 h PI) / ∑(all quantified times or conditions). Adjusted intensities were divided by the number of theoretical tryptic peptides from each protein between 7 and 30 amino acid residues in length to give an estimated IBAQ value (**Table S5**). iBAQ abundance values for viral proteins in DCs were compared to values previously generated for HFFFs^25^. Prior to calculation of the DC:HFFF iBAQ ratio, for each viral protein quantified in both cell types, iBAQ values were first normalised to the total abundance of these proteins. For example, the DC iBAQ value for UL123 was normalised to the total of DC iBAQ values for viral proteins quantified in both DCs and HFFFs. The ratio of these normalised values (DC:HFFF) was plotted in **Figure 4A**. For **Figure 4B**, to enable comparison between cell types, the relative abundance for each viral protein in DCs was first adjusted using its DC:HFFF iBAQ ratio (shown in **Figure 4A** and **Table S5**), then all values for each protein were normalised to a maximum value of 1.

Where PSMs had been assigned to a non-canonical viral ORF but were redundant to a canonical viral protein, peptides corresponding to the canonical protein were separated from those unique to the ORF, generating two separate entries as described above. For the non-canonical ORF, the number of theoretical peptides from the non-canonical protein fragment were used in the IBAQ calculation.

#### Statistical analysis

The exact value of n within figures is indicated in the respective figure legends, and refers to the number of biological replicates. Blinding or sample-size estimation was not appropriate for this study. There were no inclusion criteria and no data were excluded.

*Figures 1, 2, 3, 4, 6, S1-4*. Whole-cell and plasma membrane experiments were performed in biological duplicate. The method of significance A was used to estimate the p-value that each ratio was significantly different to 1^73^. Values were calculated and corrected for multiple hypothesis testing using the method of Benjamini-Hochberg^74^. A corrected p < 0.05 was considered statistically significant.

*Figures 3, S1, S2*. Pearson correlation values were calculated in Excel.

*Figure 4.* A one-sample Wilcoxon test was used to determine whether the log_2_ fold change of normalised iBAQ values was significantly different to 0. P-values were calculated and adjusted for multiple-hypothesis testing using the Benjamini-Hochberg method in R version 4.2.2.

*Figure 5.* GraphPad Prism 8 software (GraphPad Software, Inc., CA, USA) was used for Figure 5. Data sets with multiple samples were analysed by either one-way ANOVA test with Tukey’s post-hoc analysis (Figure 5A) or two-way ANOVA test with Sidak’s post-hoc analysis (Figure 5B and Figure 5C). All data was plotted as mean ± standard deviation (SD) unless otherwise stated. Significance was assigned as follows; *p < 0.05; **p <0.01; ***p< 0.001 and ****p<0.0001.

#### Pathway analysis

The Database for Annotation, Visualisation and Integrated Discovery (DAVID) was used to determine pathway enrichment^21,22^. A given cluster was always searched against a background of all proteins quantified within the relevant experiment.

#### Data and software availability

Unprocessed peptide data files for Figures 1, 2, 3, 4 and 6 are available at https://data.mendeley.com/preview/48w9d4z6c3?a=73bf2acf-8b84-4d68-9864-6f5933406acc.

These files include details of peptide sequence, redundancy, protein assignment raw unprocessed TMT reporter intensities and isolation specificity. The mass spectrometry proteomics data have been deposited to the ProteomeXchange Consortium (http://www.proteomexchange.org/) via the PRIDE partner repository. Reviewer account details: username, password 0v4FtR6k.

## Acknowledgements

We are grateful to Prof. Steve Gygi for providing access to the “MassPike” software pipeline for quantitative proteomics. LKJ was supported by a PhD studentship from the Systems Immunity Research Institute (SIURI) at Cardiff University, and by a KRUK project grant (RP_008_20210729) awarded to ECYW, RJS and Sian Griffin. NCH was supported by a PhD studentship from Health and Care Research Wales. This work was supported by MRC (MR/S00971X/1) and Wellcome Trust (226615/Z/22/Z) grants to RJS and ECYW, a MRC project grant (MR/V000489/1) to ECYW, DAP, and RJS, an MRC Project Grant (MR/X000516/1) to MPW, and the NIHR Cambridge Biomedical Research Centre (NIHR203312). Flow cytometry data were acquired using an Attune NxT (Thermo Fisher) which was obtained and serviced with grants from the MRC (MR/P001602/1, MR/S00971X/1, MR/V000489/1), and Wellcome Trust 204870 (P Griffiths, UCL) and 207503/Z/17/Z (I Humphreys, Cardiff University). For the purpose of open access, the authors have applied a CC BY public copyright license to any author accepted manuscript version arising from this submission.

---

**Table S9.** Thermocycling programme, Comparative CT with Melt, used for qPCR in this project.

| *Stage* | *Temperature (°C)* | *Time* | *Data Collection* | *Cycles* |
| --- | --- | --- | --- | --- |
| *Stage 1* | 50 | 2 min |  | 1 |
|  | 95 | 10 min |  |  |
| *Stage 2* | 95 | 15s |  | 40 |
|  | 60 | 1 min | ✓ |  |
| *Melt Curve* | 95 | 15s |  | - |
|  | 60 | 1 min | ✓ |  |
|  | 95 | 15s |  |  |

**Figure S1.**
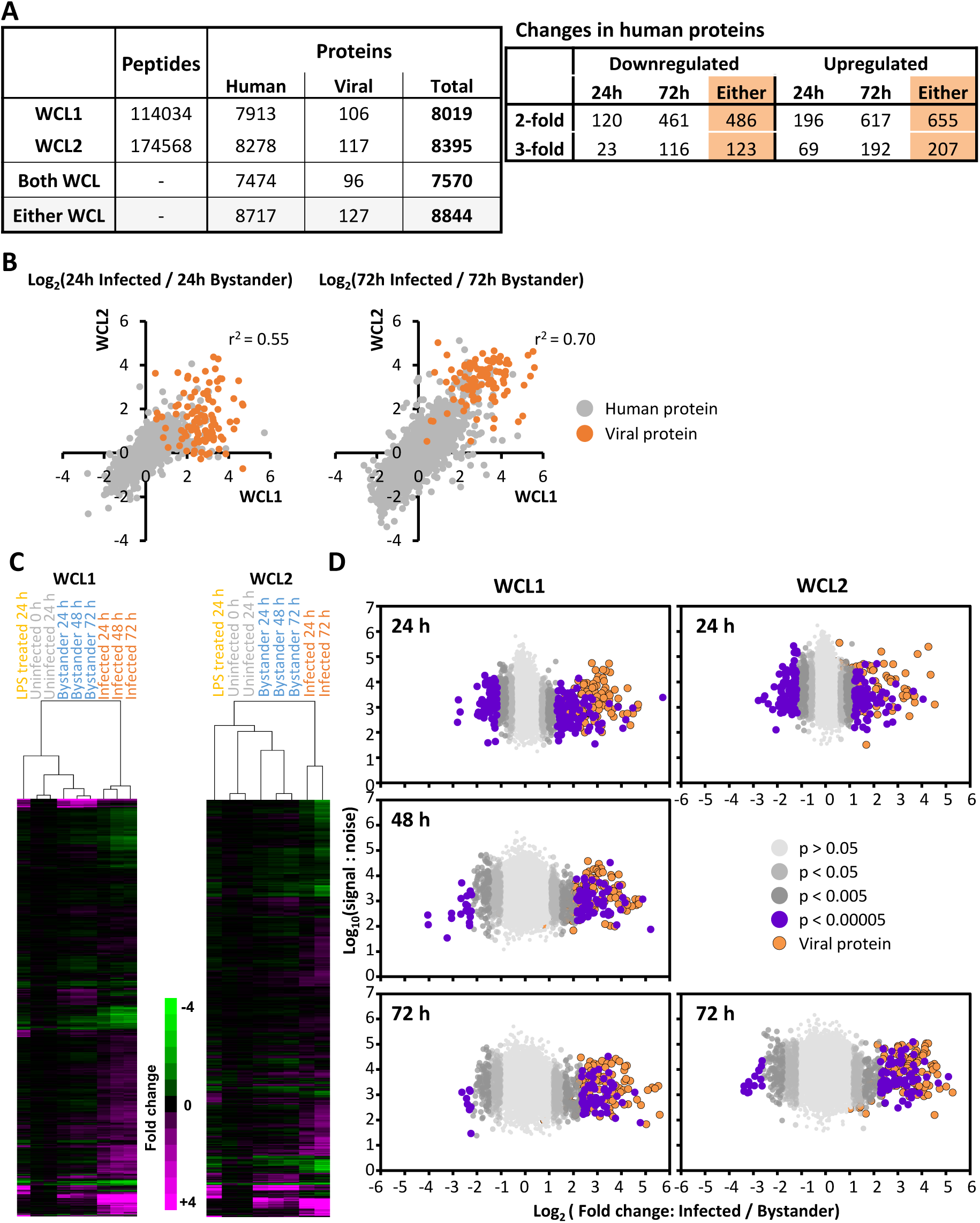
Comparison of the two whole cell lysate proteomic datasets. (A) Peptides and proteins quantified in each dataset, and regulation of human proteins at 24 or 72 hpi. Average data were used for proteins quantified in both experiments. (B) Correspondence between the two WCL datasets, comparing fold changes between infected and bystander samples. (C) Hierarchical cluster analysis of all proteins quantified in each WCL dataset. The average S:N of the two uninfected time points (0 and 24 hpi) was used for comparison with 24 and 72 hpi. Average data were used for proteins quantified in both experiments. (D) Scatterplot of all proteins quantified in experiments WCL1 and WCL2. Benjamini-Hochberg adjusted Significance A values were used to estimate p-values (see Methods). Data from 48 hpi was only available for WCL1.

**Figure S2.**
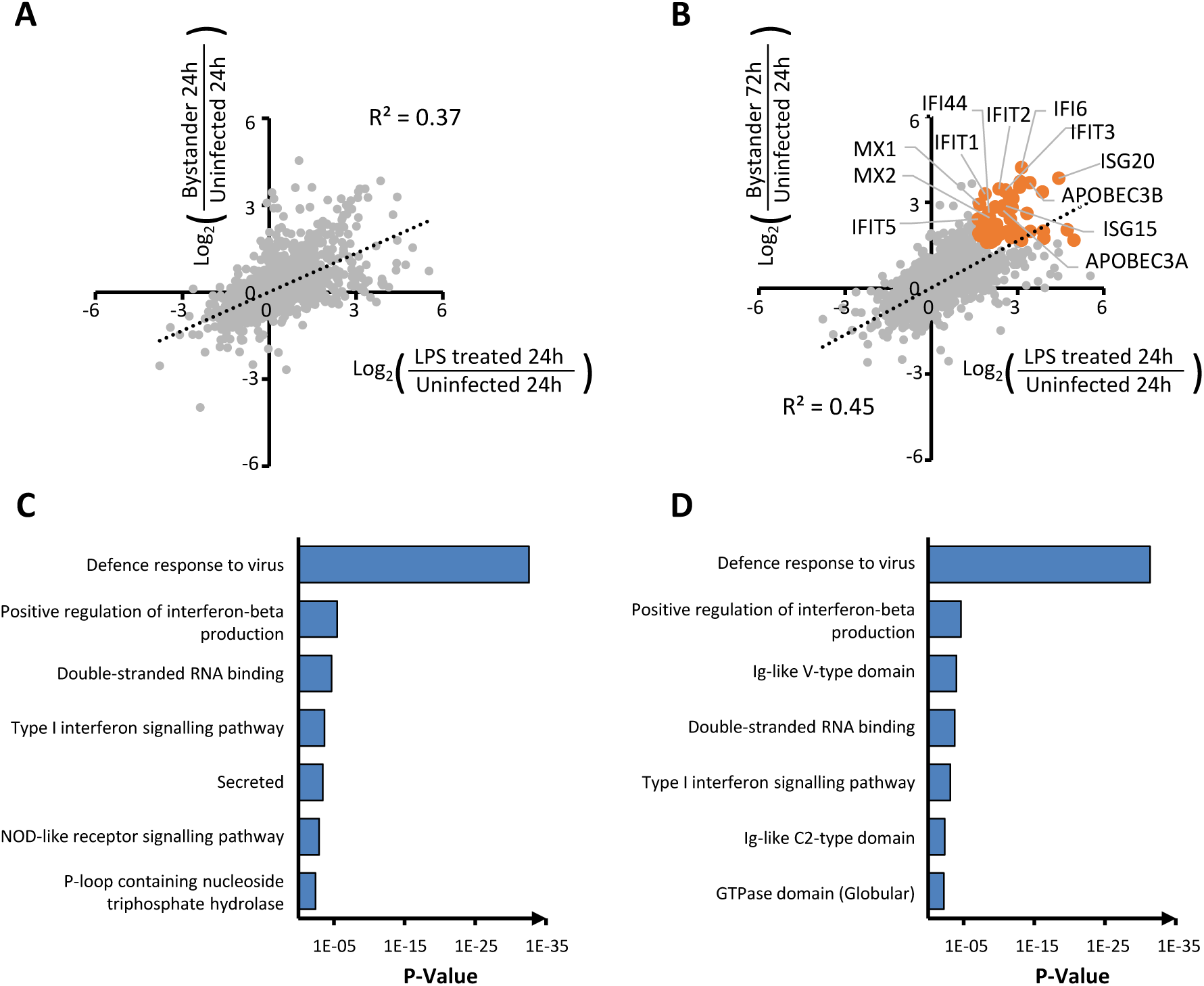
Similar proteomic changes in bystander DC and DCs stimulated with LPS. (A) Scatterplot of proteins quantified in either experiment WCL1 or WCL2, with average fold change shown where quantified in both. Changes in bystander DCs at 24 hpi correlated with changes in LPS-stimulated DCs. (B) Scatterplot as for (A) but showing changes in bystander DCs at 72 hpi. Some of the most substantially upregulated proteins (orange dots) included interferon-stimulated genes. (C-D) Functional enrichment analysis of 108 and 143 proteins upregulated >2-fold in both bystander cells and in LPS-stimulated DCs, from (A) and (B) respectively.

**Figure S3.**
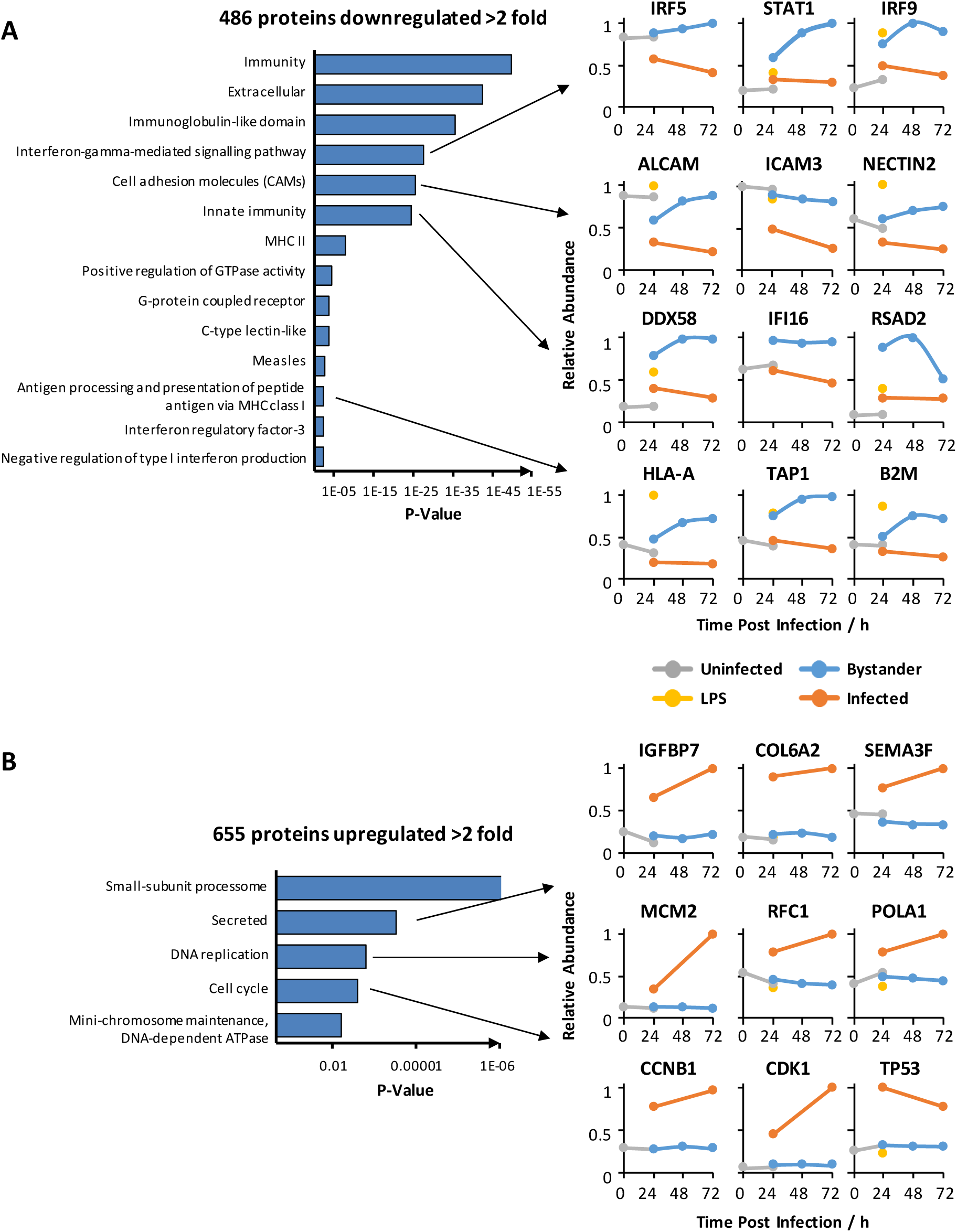
Enrichment analysis of proteins >2-fold down- or up-regulated in infected cells compared to bystander cells. (A) Functional enrichment analysis of the 486 proteins downregulated >2 fold. Displayed are significantly enriched clusters (Benjamini corrected p < 0.01) with example protein profiles. Full data for clusters enriched at p<0.05 are shown in **Table S3A**. (B) Functional enrichment analysis of the 655 proteins upregulated >2 fold. Displayed are significantly enriched cluster (Benjamini corrected p < 0.05) with example protein profiles. Full data for clusters enriched at p < 0.05 are shown in **Table S3B**.

**Figure S4.**
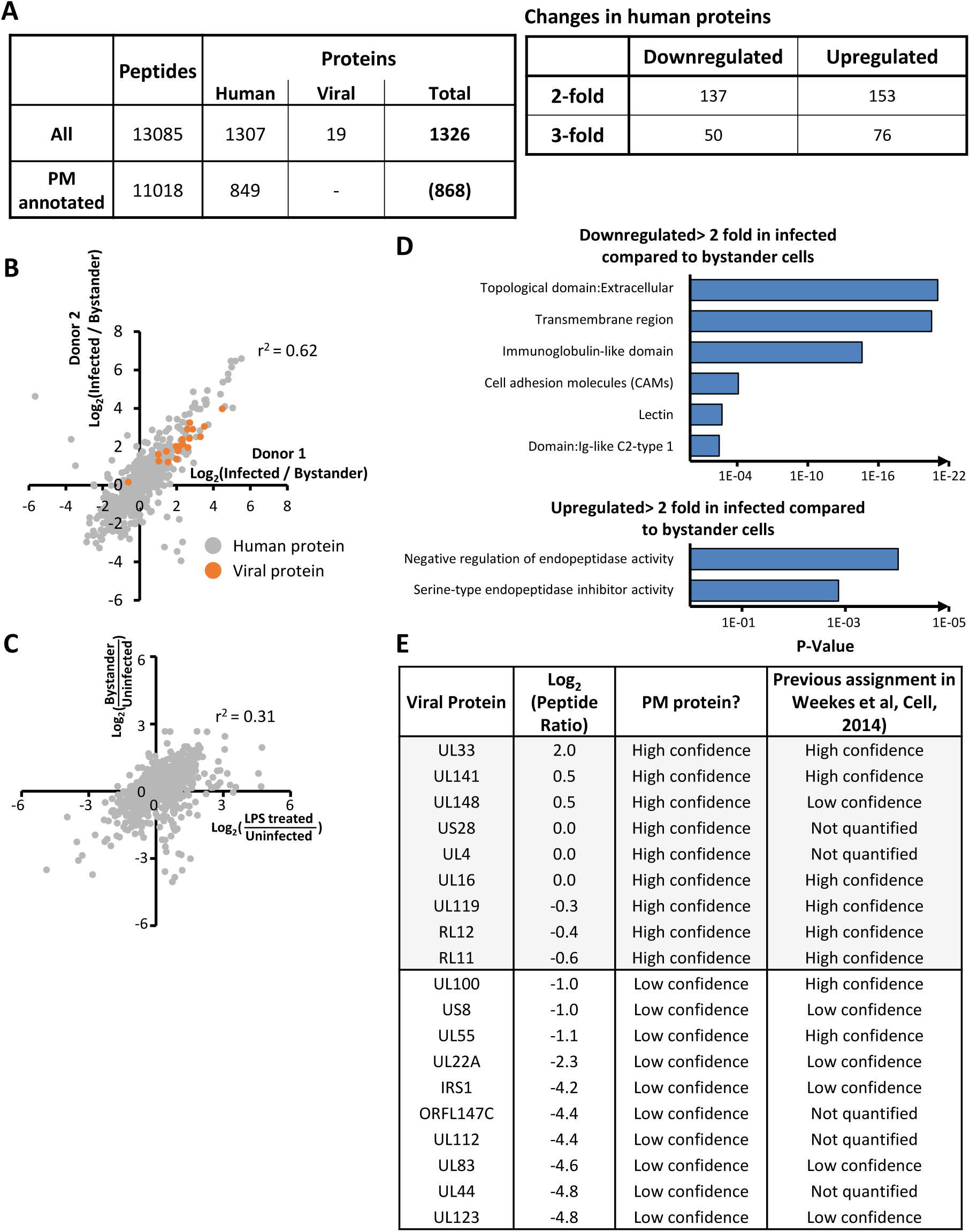
Comparison between and further analysis of the two PM proteomic datasets. (A) Peptides and proteins quantified, and regulation of human proteins in infected DCs. Fold changes were calculated from the average of the two replicate samples. All proteins were quantified in both experiments, as part of the same TMT multiplex. (B) Scatterplot showing correspondence between the two PM datasets, comparing fold changes between infected and bystander samples. (C) Scatterplot showing correspondence between average fold changes in PM proteins from bystander DCs and DCs stimulated with LPS. (D) Functional enrichment analysis of the proteins regulated >2 fold. Displayed are significantly enriched clusters (Benjamini corrected p < 0.01) with example protein profiles. Full data for clusters enriched at p < 0.05 are shown in **Table S6**. (E) Viral proteins present at the DC cell surface with (log_2_(peptide ratio) > -1.0, see **Figure 6D**). Confidence assignments from HFFF data in Weekes et al 2014 are shown for comparison.

